# Acute silencing uncovers multiple forms of activity-dependent neuronal survival in the mature entorhinal cortex

**DOI:** 10.1101/2021.10.18.464827

**Authors:** Rong Zhao, Stacy D. Grunke, Ming-Hua Li, Caleb A. Wood, Gabriella A. Perez, Melissa Comstock, Anand K. Singh, Kyung-Won Park, Joanna L. Jankowsky

## Abstract

Neurodegenerative diseases are characterized by selective vulnerability of distinct cell populations; however, the cause for this specificity remains elusive. Many circuits that degenerate in disease are shaped by neural activity during development, raising the possibility that mechanisms governing early cell loss may be misused when activity is compromised in the mature brain. Here we show that electrical activity and synaptic transmission are both required for neuronal survival in the adult entorhinal cortex, but these silencing methods trigger distinct means of degeneration in the same neuronal population. Competition between active and inactive cells drives axonal disintegration caused by synaptic inhibition, but not axon retraction due to electrical suppression. These findings suggest that activity-dependence may persist in some areas of the adult brain long after developmental critical periods have closed. We speculate that lifelong plasticity required to support memory may render entorhinal neurons vulnerable to prolonged activity changes in disease.

## Introduction

Each neurodegenerative disease is characterized by selective loss of a specific neuronal population. Current research efforts are focused on the molecular features of these neurons that render them susceptible to disease pathology. Modern transcriptomic techniques have revealed distinct gene signatures of vulnerable neuronal populations, but this approach takes a static view of circuits that remain active throughout life. Neuronal activity plays a key role in circuit refinement during development up to and including cell death, raising the possibility that developmental mechanisms may be aberrantly reactivated in the context of disease. The convergence of development and degeneration has been described previously with the identification of common kinase signaling pathways and cell cycling markers (Bothwell and Giniger, 2000; Nguyen et al., 2002). However, these studies overlook the contribution of dynamic, circuit-level features that initiate intracellular signaling pathways to stimulate cell death in vulnerable neurons. During CNS development, some regions eliminate up to 50% of immature neurons through activity-dependent programmed cell death (Murase et al., 2011; Wong and Marin, 2019). Still more complex and varied processes drive axonal refinement in development (Neukomm and Freeman, 2014; Riccomagno and Kolodkin, 2015). Despite being one of the strongest factors in developmental circuit refinement, neuronal activity has not been well studied as a potential contributor to neurodegeneration (Gonzalez-Rodriguez et al., 2020; Wang and Holtzman, 2020).

Developmental circuit refinement is historically characterized by critical windows which are transient periods of heightened plasticity when synapse pruning, axon elimination, and cell death are governed by sensory experience. Many sensory systems use this critical period to establish precisely mapped connections between external input and cortical target areas (Fox and Wong, 2005; Hensch and Fagiolini, 2005; Reh et al., 2020). Competition is a common theme in determining which neural connections persist and which are eliminated (Blakemore, 1991; Zhao and Reed, 2001; Yu et al., 2004; Fox and Wong, 2005; Hensch and Fagiolini, 2005; Yasuda et al., 2011). Manipulations to sparsely suppress or enhance activity in developing circuits can have dramatic effects on the final pattern of innervation. These manipulations are most effective during the postnatal period, but recent work reveals that the traditional concept of a transient window for circuit refinement may be overly restrictive. Experimental manipulation of innate activity uncovered latent reorganization in the mature visual and auditory systems, while artificial electrical stimulation of the adult corticospinal tract can partially reverse deficits caused by developmental deprivation (Salimi et al., 2008; Bavelier et al., 2010; de Villers-Sidani and Merzenich, 2011; Hubener and Bonhoeffer, 2014; Nahmani and Turrigiano, 2014; Martin, 2016). The manipulations required to evoke wholesale structural changes in the mature brain are often more extreme than those required in young animals, but in some contexts the outcome for circuit reorganization is similar (de Villers-Sidani and Merzenich, 2011; Hubener and Bonhoeffer, 2014; Nahmani and Turrigiano, 2014).

This newly appreciated potential for circuit reorganization in the mature brain might also contribute to neurodegeneration in disease. Changes in neuronal activity are an early hallmark of several degenerative disorders, and it is increasingly clear that network dysfunction can drive pathogenesis (Cirrito et al., 2005; Yamada et al., 2014; Yamamoto et al., 2015; Palop and Mucke, 2016; Wu et al., 2016; Yuan and Grutzendler, 2016; Harris et al., 2020; Rodriguez et al., 2020). Admittedly, most activity-dependent structural changes described in the mature brain focus on shifts in innervation rather than the overt cell death of development and neurodegeneration. Additionally, sensory areas where circuit reorganization has been best documented are generally not affected in neurodegeneration. We were therefore surprised to uncover evidence for activity-dependent cell death mechanisms in a chemogenetic model of circuit dysfunction in the adult entorhinal cortex (EC). EC neurons are among the earliest site of neuropathology in AD and loss of these cells impairs the main conduit of information flow into the hippocampus (Hyman et al., 1984; Van Hoesen et al., 1986, 1991; Gomez-Isla et al., 1996; Kordower et al., 2001). While modeling the circuit-level consequences of this early degeneration, we uncovered evidence that the EC maintains activity-dependent cell death mechanisms well into adulthood. Using multiple chemogenetic systems, we show that different forms of neuronal silencing elicit distinct patterns of axon degeneration prior to neuronal cell loss. One of these manipulations is governed by classic inter-neuronal competition, while another appears to be cell-autonomous. Together, our studies indicate that neuronal activity plays an ongoing role in the survival of EC neurons and raise the possibility that perturbations in activity may partner with molecular pathology to drive degeneration of this region.

## Results

### Unexpected loss of EC2 neurons following chloride-based neuronal silencing

In previous work we described a chemogenetic model for studying neuronal dysfunction in the adult EC (Zhao et al., 2016). This model used an engineered chloride channel to allow ligand-activated disruption of action potential initiation in a subset of EC layer 2 neurons (EC2) (Lynagh and Lynch, 2010, 2012). The modified channel (GlyCl) was expressed in EC2 neurons using a tetracycline-transactivator driver line that was selective for, but not exclusive to, these cells (Nop-tTA; TRE-GlyCl-YFP) (Yasuda and Mayford, 2006; Yetman et al., 2015). The GlyCl channel is activated by the ligand ivermectin (IVM) and was co-expressed with yellow fluorescent protein (YFP) to identify the silenced cells. Bath application of 100 nM IVM to acute brain slices suppressed neuronal firing by up to 90% in YFP+ entorhinal neurons from GlyCl transgenic mice, but had no effect on EC neurons from control animals (Figure 1A). We previously demonstrated that systemic administration of IVM decreased neuronal firing in vivo for several hours after intraperitoneal injection and that the drug half-life of >12 hours likely maintained this effect for a day or more (Zhao et al., 2016).

**Figure 1.**
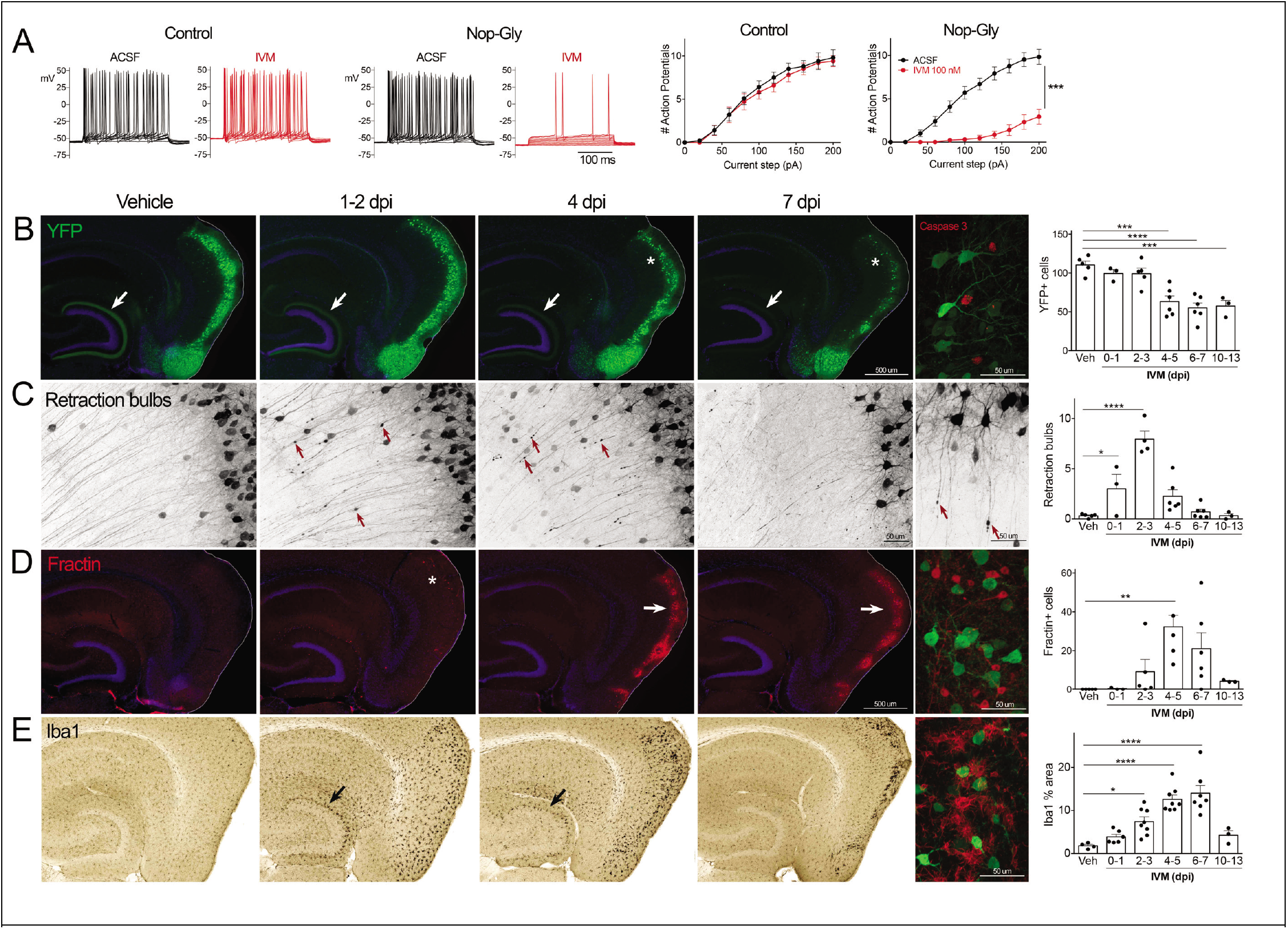
Transient electrical silencing with GlyCl causes cell death in EC neurons. A. Example traces from current-clamp recordings in brain slices from control and GlyCl-expressing mice, with and without 100 nM IVM. Graphs on right show average number of action potentials elicited at each current step. Data expressed as mean ± SEM, two-way repeated measure ANOVA (p value shown for interaction of current x treatment). n=10 cells from 10 slices (5 mice, control WT or single transgenic); 13 cells from 13 slices (6 mice, Nop-tTA; TRE-GlyCl-YFP bigenic). B. YFP immunostaining reveals that silencing EC neurons lead to progressive loss of dentate innervation (white arrow) followed by loss of YFP-labeled neuronal cell bodies in the EC (asterisk). Final image shows the presence of active caspase-3 immunostaining (red) among YFP+ cell bodies (green) at 3 days after IVM injection (dpi). C. High magnification images in EC show retraction bulbs on YFP-labeled axons withdrawing from the DG (red arrows). Final image shows single YFP-labeled axons terminating in retraction bulbs at 2 dpi. D. Fractin immunostaining marks the emergence of damaged/dying neurons in EC. Final image shows fractin+ cells (red) among YFP+ cells at 7 dpi. E. Iba1 immunostaining reveals early microglial activation upon denervation of the DG (black arrow) followed by pronounced reactivity in the superficial EC. Final image shows Iba1 (red) and YFP (green). Graphs show average values per section for each animal ± SEM. n=4-5 (veh), 3-6 (0-1 dpi), 4-8 (2-3 dpi), 4-8 (4-5 dpi), 6-7 (6-7 dpi), 3 (10-13 dpi). All comparisons were done by one-way ANOVA with Dunnett’s post-hoc test, *p<0.05, **p<0.01, ***p<0.001, ****p<0.0001.

Following the initial description of this new chemogenetic silencing system, we discovered an unexpected change in the labeling of GlyCl+ EC neurons over the week following IVM injection. Vehicle injected animals displayed clear YFP labeling of neuronal cell bodies in EC2 and their axonal projections into the dentate gyrus (DG) (Figure 1B). In contrast, animals injected with a single dose of IVM (5 mg/kg, i.p.) showed diminished DG axonal labeling within 1-2 days post injection (dpi) and labeling was nearly undetectable by 4 dpi. Alongside this change in the DG, the number of labeled cell bodies in EC2 was significantly decreased by 4 dpi and reduced by 50% at 7 dpi (Figure 1B).

The spatiotemporal pattern of diminished labeling suggested that the silenced cells may have been eliminated from the circuit. Four pieces of evidence supported the hypothesis that neurons had died after silencing rather than simply shutting down expression of the fluorescent protein. First, we found EC cells labeled with active caspase-3, indicative of apoptotic cell death, in the days immediately following IVM injection (Figure 1B). Caspase-labeled cells were sporadic, but only observed in EC2 of animals subjected to entorhinal silencing. Second, many YFP-labeled EC2 axons developed retraction bulbs, which are typically observed after axonal injury (Hill et al., 2016) (Figure 1C). These bulbs appeared within 1 day of IVM treatment, peaked at 2-3 dpi, and were gone over the same time course as the loss of YFP-labeled cells (Figure 1C). Third, EC2 cells contained a caspase-cleaved actin fragment (fractin), again consistent with apoptotic cell death (Yang et al., 1998; Schulz et al., 2009) (Figure 1D). Fractin labeling peaked between 4-7 days after IVM and then abated. Fourth, we observed hypertrophic Iba1+ microglia in both the DG and the EC. Iba1 staining increased by 2 days after IVM, peaked by 4-7 dpi, and receded by 10-13 dpi (Figure 1D). Collectively, our data suggests that transient neuronal silencing of EC2 neurons initiated a cascade of events beginning with axonal retraction and ending in caspase-associated cell death with a strong but focal microglial response.

### GlyCl-based cell death is largely restricted to EC2

The loss of EC2 neurons after GlyCl silencing was unexpected and initially caused us to suspect that we had simply generated a chemogenetic system for cell death. We therefore tested the effect of GlyCl silencing in other neuronal populations. The neuropsin-tTA driver line used to express GlyCl in EC2 is also active in connected regions of the retrosplenial cortex, presubiculum, and parasubiculum, which allowed us to test whether silencing elicited cell death outside of EC2 (Yetman et al., 2015) (Figure 2A-C). We first confirmed that GlyCl expression was sufficient to suppress neuronal firing upon IVM exposure in these additional populations. Action potential firing was significantly reduced in YFP+ neurons from acute brain slices for all three areas upon bath application of 100 nM IVM (Figure 2D-G and Supplemental Figure 1). Action potential firing was somewhat better repressed in EC2 than in parasubiculum at higher current steps, followed by presubiculum and retrosplenial cortex, in a pattern that was consistent with the relative intensity of YFP fluorescence in each of these areas (data not shown). We counted GlyCl+ cells in each brain area before and after systemic IVM treatment at 5 mg/kg. To aid cell counting, we added a third transgene to co-express nuclear-localized lacZ in GlyCl+ cells (Nop-tTA; TRE-GlyCl-YFP; TetO-nls-lacZ-GFP). We began by confirming that EC2 cells were still diminished by GlyCl silencing in mice carrying this added nuclear lacZ transgene. As before, a single dose of IVM significantly reduced the number of labeled EC2 cells in GlyCl+ mice (Figure 2H). Because the breeding also generated animals expressing entorhinal lacZ without GlyCl, we were further able to show that IVM treatment has no effect on neuronal survival in the absence of the channel (Figure 2I). Importantly, we found no cell loss in any brain region other than EC2 in mice that co-expressed lacZ with GlyCl (Figure 2J-K). Fractin immunostaining and changes in Iba1 density were occasionally found in the parasubiculum bordering EC2, but not in the remainder of parasubiculum, presubiculum, or retrosplenial cortex (data not shown).

**Figure 2.**
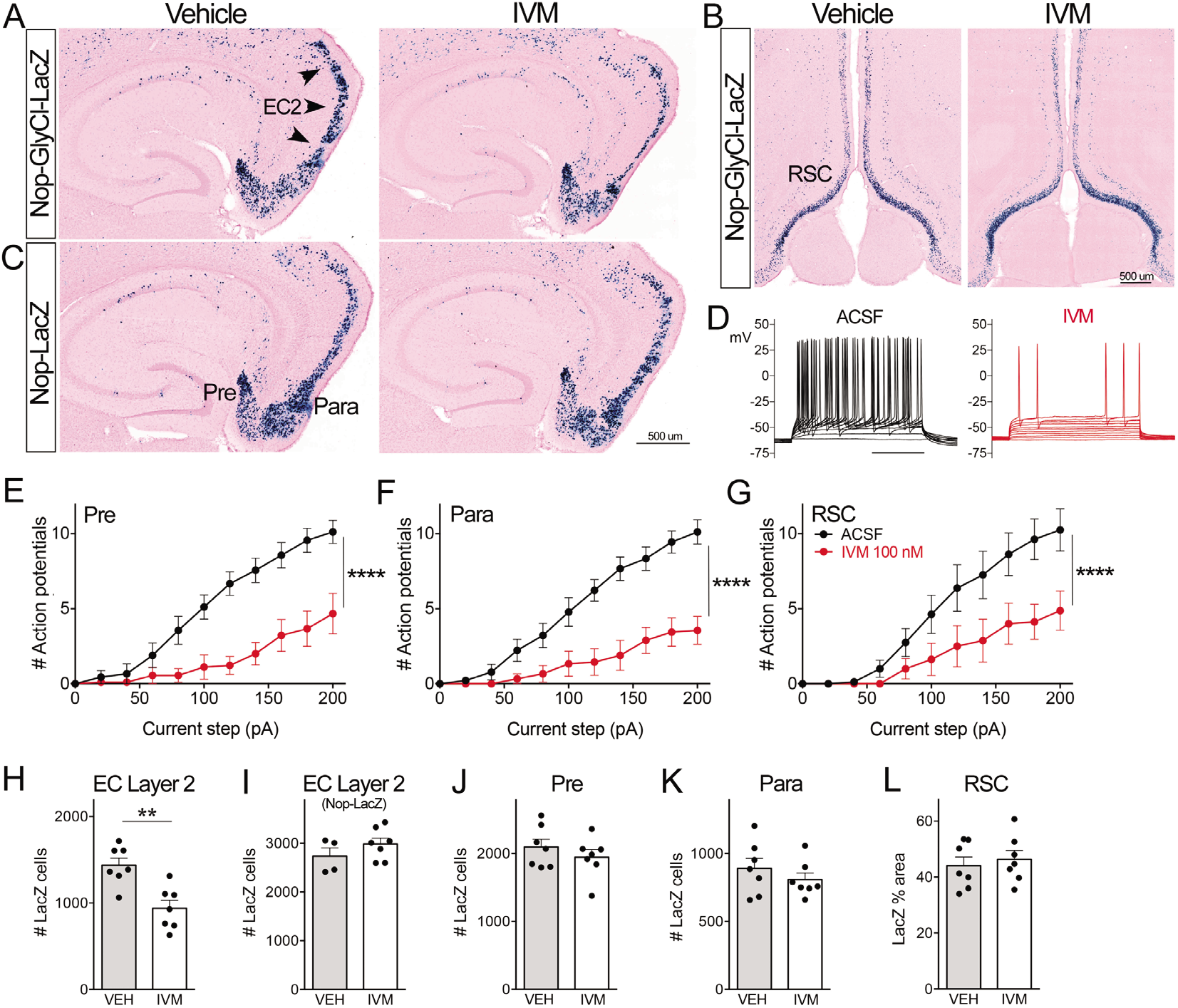
Electrical inactivation of other brain regions does not cause neuronal loss in the Nop-GlyCl model. A-B. The Nop-tTA line drives transgenic GlyCl expression in cells beyond the EC, including pre- and parasubiculum and retrosplenial cortex (RSC). Co-expressed nuclear lacZ was used to identify GlyCl+ cells in mice with both transgenes. C. Animals expressing lacZ without GlyCl were used to control for any cell death caused by IVM itself. D. Example traces from a presubiculum neuron recorded by current-clamp in an acute brain slice before and after IVM application. E-G. Whole-cell recordings of presubiculum (E), parasubiculum (F), and retrosplenial cortex (G) show effective suppression of action potential firing upon IVM perfusion in each area. Data expressed as mean ± SEM, one-way ANOVA with Dunnett’s post-hoc test (two sided). Pre n=9 cells from 9 slices (4 mice); para n=9 cells from 9 slices (4 mice); RSC n=8 cells from 8 slices (6 mice). H-L. The number of lacZ+ cells remaining in EC2 is diminished by IVM in mice that express GlyCl (H) but not in mice that lack the channel (I). No other area shows cell loss following GlyCl neuronal silencing: presubiculum (J), parasubiculum (K), retrosplenial cortex (L). Cell counts were done 30 dpi. Graph data expressed as summed cell counts (EC, Pr/Para), or average % area (RSC) for each animal ± SEM, Student’s t-test (two sided). EC2+GlyCl n=7 veh, 7 IVM; EC2 lacZ only n=4 veh, 7 IVM; pre, para, RSC n=7 veh, 7 IVM. **p<0.01

### Other means of silencing via ion flux elicit degeneration of the perforant path

We next sought to determine whether EC2 neurons were selectively sensitive to changes in chloride flux induced by the GlyCl system or if instead they were more broadly vulnerable to electrical silencing. We tested two means of electrical inactivation dependent on potassium rather than chloride. Our first experiments used a mutated Kir 2.1 channel to suppress firing by potassium shunting (Xue et al., 2014). To express the Kir channel in EC2, we stereotaxically injected a tTA-dependent Kir2.1-YFP viral construct into the EC of Nop-tTA single transgenic mice. Viral expression peaked at approximately 7 dpi, when field recordings of input-output responses in the perforant path demonstrated a significant reduction in transmission between EC2 and DG (Supplemental Figure 2). Co-expressed YFP again allowed us to visualize the loss of axonal labeling from the DG over the following weeks, qualitatively similar to but slower than what we observed with GlyCl silencing (Supplemental Figure 2). Iba1 immunostaining increased to a peak between 13-16 dpi in both EC and DG of Kir-expressing animals, consistent with neuronal damage in these areas. Control animals received TRE-YFP virus without Kir and showed no loss of fluorescence nor any change in Iba1 beyond the needle track in either the DG or EC over 4 weeks post injection (Figure 3).

**Figure 3.**
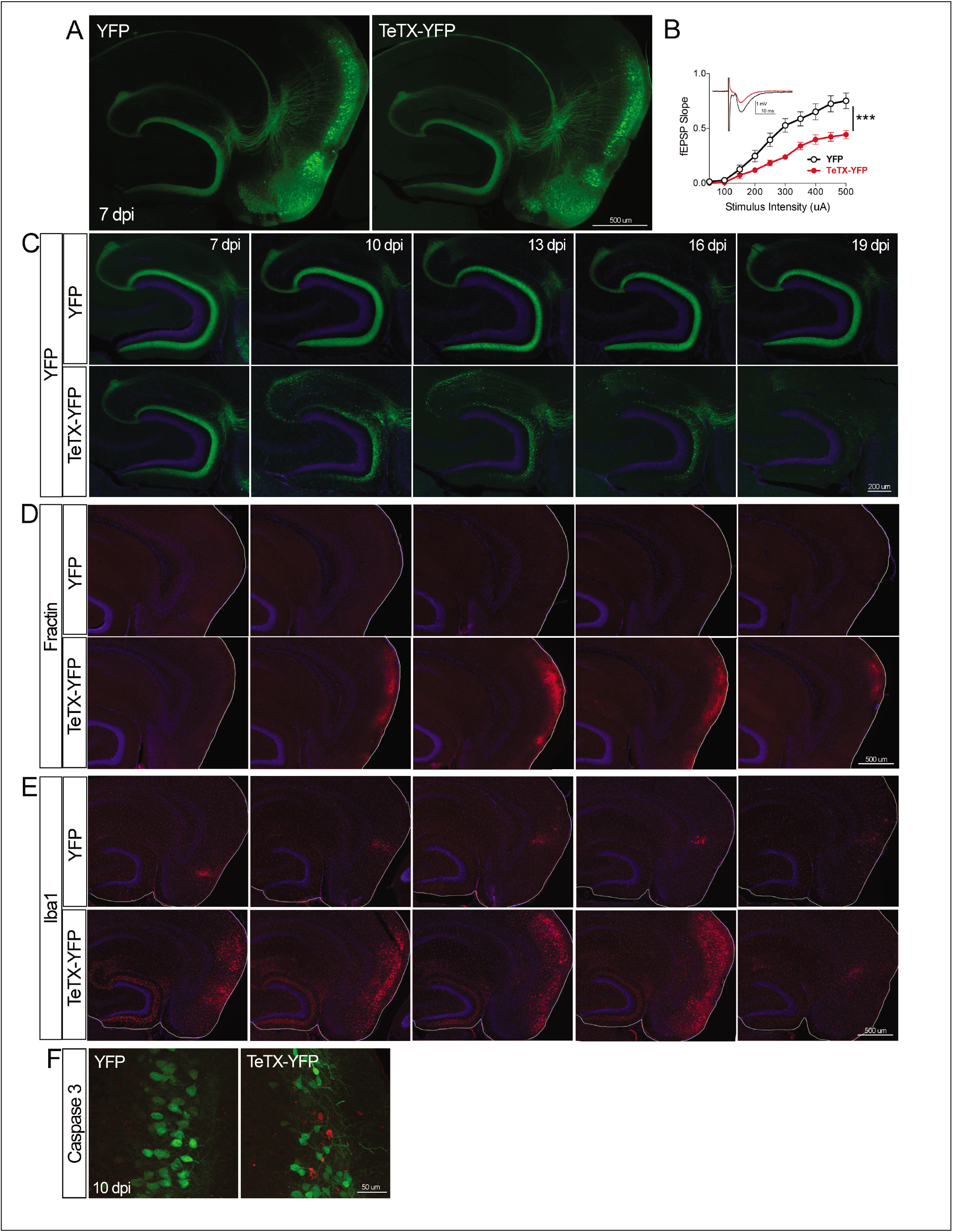
Entorhinal cell death is also elicited by synaptic silencing with tetanus toxin. A. AAV was used to express either YFP or YFP plus TeTX in entorhinal neurons. By 7 dpi, axons terminating in the DG can be clearly seen in both constructs by native fluorescence. B. Virally-delivered TeTX effectively diminishes synaptic transmission into DG. Data expressed as mean ± SEM, two-way repeated measure ANOVA (p value for genotype shown, ***p<0.001). n=15 slices from 3 mice (YFP), 17 slices from 4 mice (TeTX), harvested 7-9 dpi. Note that YFP data shown here is identical to Supplemental Figure 2B as the same control group was used for both experiments. C. TeTX causes disintegration of labeled perforant path axons. No change in dentate innervation is seen in animals injected with YFP control virus. DAPI counterstain indicates the dentate granule cell layer. D. Fractin immunostaining appears in EC as axons disintegrate in DG. E. Microglial activation detected with Iba1 appears rapidly in both DG and EC by 7 dpi in mice injected with AAV-TeTX and persists in EC for more than a week before resolving. Microglial activation is seen only at the injection site in mice injected with AAV-YFP. F. Active caspase 3 immunostaining (red) is found among YFP+ cells in the EC (green) only in animals co-expressing TeTX. 4 dpi n=2 YFP, 2 TeTX; 7 dpi n=4 YFP, 4 TeTX; 10 dpi n=3 YFP, 4 TeTX; 13 dpi n=5 YFP, 4 TeTX; 16 dpi n=6 YFP, 4 TeTX; 19 dpi n=3 YFP, 5 TeTX; 22 dpi n=2 YFP.

We also suppressed firing of EC2 neurons using the inhibitory DREADD hM4Di, which functions through G-protein coupling to Kir channels (Armbruster et al., 2007). We injected a tTA-dependent hM4Di-YFP virus into neonatal Nop-tTA mice to elicit broad DREADD expression in the EC of adult mice. The DREADD system is most often used for its short-acting effect following systemic administration of its ligand clozapine-N-oxide (CNO). In our case, reproducing the effect of GlyCl inactivation required a longer period of suppression, which we achieved by continuous CNO infusion via subcutaneous osmotic pump. Adult animals were treated with CNO or saline for 3 days and harvested 7 days after drug onset. As with GlyCl-based silencing, there was a marked loss of YFP labeling in the DG of CNO-treated mice, accompanied by the appearance of reactive microgliosis in both the DG and EC (Supplemental Figure 3). We note that we also tried this experiment using a tTA-dependent hM4Di transgenic line mated with the Nop-tTA driver (Zhu et al., 2014). Bigenic adult offspring were treated for 3 days with CNO or saline, but showed no change in labeling between conditions (data not shown). We suspect that viral hM4Di delivery may have succeeded in replicating the Kir and GlyCl findings by achieving higher expression levels and more complete or persistent silencing than the transgenic system. In both the transgenic and viral delivery methods, we were asking the DREADD system to provide prolonged suppression for which it was not designed. Taken together, the Kir and viral DREADD data further support the persistence of activity-dependent cell survival in the adult EC.

### EC2 neurons are also vulnerable to synaptic silencing

Neuronal silencing with GlyCl, Kir2.1, or hM4Di affects both action potential initiation and synaptic transmission. We next wanted to tease apart these effects using another chemogenetic tool that blocks one but not the other. We used virally-expressed tetanus toxin (TeTX) to suppress neurotransmitter release from EC2 neurons in adult Nop-tTA mice without affecting electrical activity. Co-expressed YFP was used to visualize the silenced neurons and YFP alone was injected as a control (Figure 3A). Field recording in acute slices from virally-injected mice confirmed that TeTX expression significantly reduced electrical transmission between EC2 and DG and that the extent of suppression was consistent with the percentage of neurons expressing the virus (Figure 3B). Expression of TeTX-YFP initially labeled both EC2 neuronal cell bodies and their axon terminals in the DG, similar to the pattern in YFP injected control mice. Viral expression peaked at 7 dpi and shortly afterwards YFP intensity began to fade in the DG of TeTX mice (Figure 3C). Loss of perforant path labeling progressed over the next 2 weeks until nearly undetectable. Animals injected with YFP alone showed no change in labeling over the same time period. Loss of axonal labeling in TeTX mice was accompanied by the emergence of fractin staining in the EC (Figure 3D). Again, the change was specific to mice expressing TeTX and was not caused by viral expression of YFP alone. Fractin-positive cells first appeared at 10 dpi, became more prominent at 13 dpi, and largely abated by 19 dpi. As with GlyCl-based silencing, we observed reactive microglia in the DG and EC as an early marker of impending cell loss (Figure 3E), and found sporadic cells labeled with active caspase 3 (Figure 3F). This cascade of events leading from impaired neurotransmission to cell death was comparable across the various silencing methods we tested, suggesting that EC2 neurons are broadly susceptible to inactivity.

### TeTX and GlyCl elicit distinct cell death processes

In all cases of EC2 silencing, we observed the loss of DG innervation over time. Closer inspection of the GlyCl and TeTX-expressing mice revealed a more nuanced story. Although electrical suppression by GlyCl caused gradual loss of DG fluorescence, synaptic inhibition by TeTX caused pronounced axonal disintegration (Figure 4). GlyCl silencing was characterized by the appearance of retraction bulbs in deeper layers of the EC, suggesting neuronal loss followed intact axonal withdraw. No retraction bulbs were visible in TeTX treated animals; instead the axonal processes became fragmented in the DG with small bright puncta that were reminiscent of Wallerian degeneration. This qualitative distinction warranted further investigation of the underlying mechanistic difference in the process of neurodegeneration caused by silencing at the cell body vs. the synapse.

**Figure 4.**
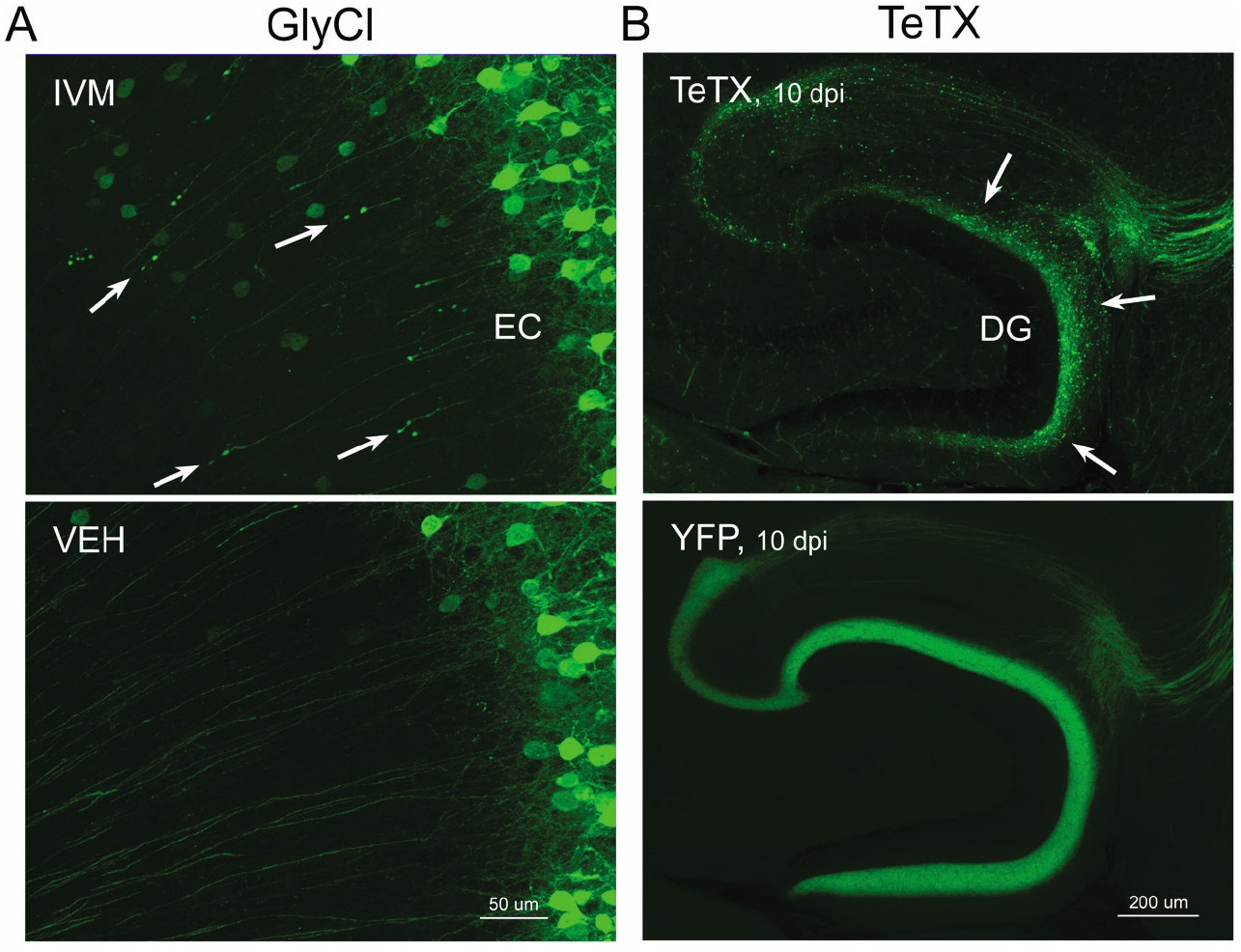
Different patterns of axonal damage attend neuronal silencing by GlyCl and TeTX. A. Retraction bulbs were visible in deep EC layers of Nop-GlyCl bigenic animals 4 days after injection of IVM (upper panel), while axons in vehicle-treated mice remain intact (lower panel). B. In contrast to the discrete retraction bulbs seen with GlyCl, TeTX silencing caused axonal fragmentation throughout the DG molecular layer (arrows, upper panel). YFP labeling in TeTX mice became punctate and innervation deteriorated within 10 days after viral injection, while the axons of EC neurons labeled with YFP alone remained robust and bright (lower panel).

### Contribution of neuronal competition distinguishes cell loss due to GlyCl from that due to TeTX

One important feature of the models described so far is that they allowed us to manipulate a targeted subset of EC2 neurons. The Nop-tTA driver line used to express GlyCl, Kir2.1, hM4Di, and TeTX is active in approximately 40% of EC2 neurons (Yasuda and Mayford, 2006). The remaining EC2 neurons were unaffected by our genetic and viral manipulations. As a result of this incomplete penetrance, we set up a situation whereby some neurons continued to fire normally while others were inactivated. Such activity-dependent competition shapes maturation of the perforant path connecting EC2 to the DG during postnatal development (Yasuda et al., 2011), and we wondered whether competition might account for neuronal loss upon silencing in the adult EC. To test this hypothesis, we chronically infused tetrodotoxin (TTX) or saline into the subiculum of Nop-GlyCl mice using a unilateral cannula fed by a subcutaneous osmotic pump. By injecting into the subiculum, the EC and DG were spared from injection site damage, but TTX would still reach the neighboring EC. The spread of TTX and efficacy of EC silencing were verified by the loss of c-fos labeling following chemically-induced seizures (data not shown). We began TTX infusion 12 hr before injecting IVM to ensure TTX had reached the EC at the time of GlyCl activation. TTX infusion was continued for another 2.5 dpi after IVM injection to suppress neuronal firing throughout the period when GlyCl might remain active (Zhao et al., 2016). Animals were euthanized 7 days after IVM to examine whether widespread inactivity caused by TTX would prevent cell death in the subset of EC2 neurons that were silenced with GlyCl.

Contrary to our hypothesis, infusion of TTX had no effect on neuronal survival in the GlyCl model. We found no difference in the area of YFP fluorescence as a surrogate for EC cell number between mice infused with TTX and saline (Figure 5A). Both IVM-treated groups lost ~66-75% of the YFP+ area compared to vehicle injected mice, similar to EC cell counts in IVM-treated GlyCl mice without cannulation (see Figure 1B). TTX infusion also had no effect on fractin expression or microglial activation evoked by IVM (Figure 5B-C). Importantly, vehicle injected mice showed no difference in YFP area between TTX and saline, indicating that TTX infusion by itself does not cause cell death. Given this negative outcome, we tested two modifications of the infusion procedure to rule out the possibility of incorrect TTX targeting or insufficient TTX duration. Neither 3 days of TTX infusion targeting the DG nor 7 days of subicular infusion changed the outcome (data not shown). These findings indicate that cell death in the GlyCl model cannot be explained by competition between the silenced cells and their active neighbors.

**Figure 5.**
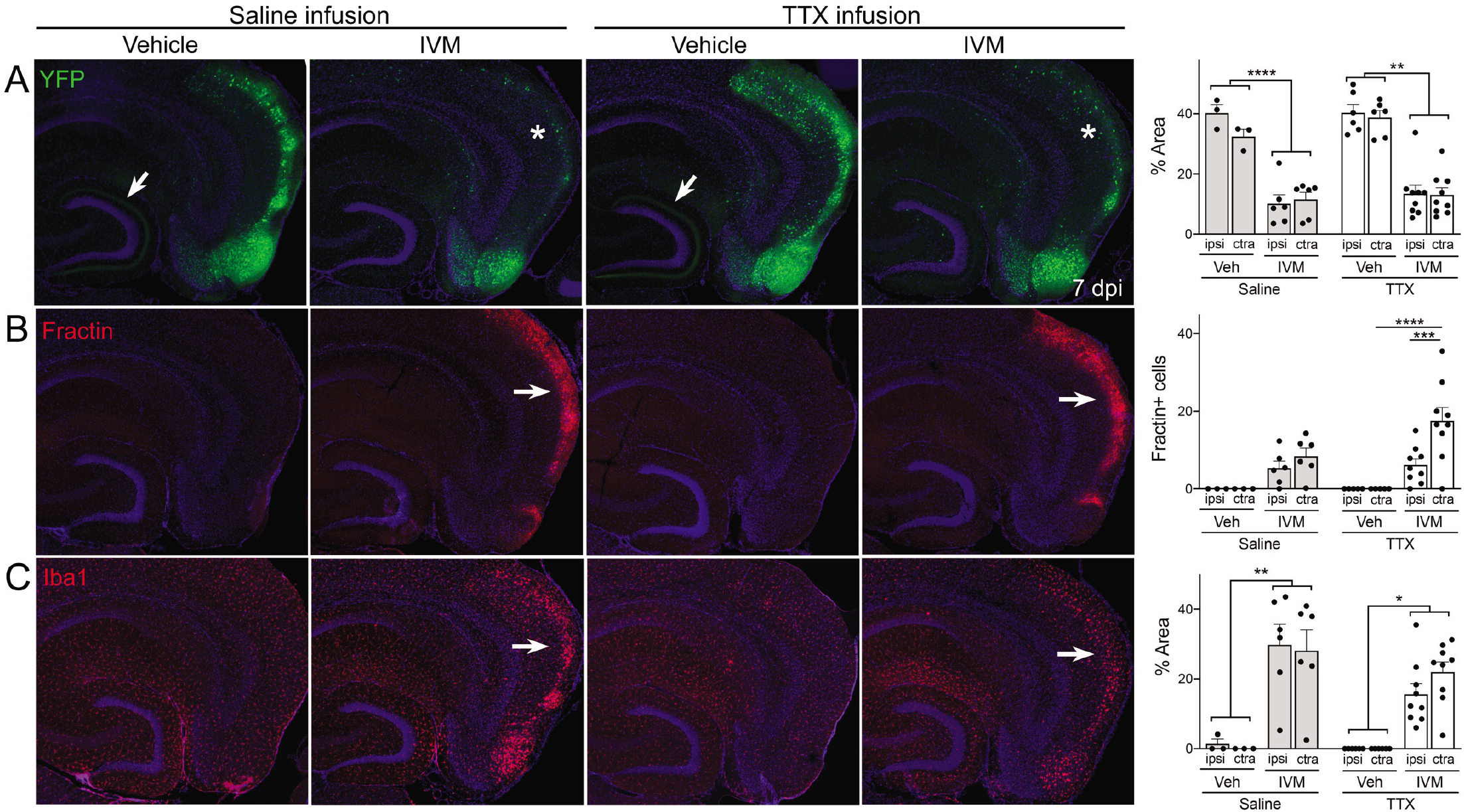
Global suppression of EC activity does not prevent cell death by GlyCl activation. Nop-tTA; TRE-GlyCl-YFP bigenic mice were cannulated to infuse either TTX or saline into the right subiculum via osmotic pump. 12 hr later, animals were injected with either vehicle or IVM. All images are from mice euthanized 7 days after IVM treatment. A. YFP immunostaining shows clear EC2 cell loss in IVM-treated mice regardless of whether they were infused with TTX or saline (asterisks). In contrast, both EC2 labeling of cell bodies and axonal labeling in the DG (arrows) were preserved in vehicle-injected mice and this pattern was unaffected by TTX infusion. B. Fractin expression was increased in EC2 of IVM-treated mice (arrows) and unchanged by TTX infusion. C. Iba1 immunostaining identified EC microglial activation in IVM-treated mice. Graph data expressed as average ± SEM, three-way ANOVA with Sidak’s post-hoc test. n=3 Vehicle+Saline infusion; n=6 IVM+Saline infusion; n=6 Vehicle+TTX infusion; n=9 IVM+TTX infusion. *p<0.05, **p<0.01, ***p<0.001, ****p<0.0001.

Due to its distinct mode of axon elimination (see Figure 4), we next tested whether neuronal competition plays a role the TeTX model. We started TTX infusion into the subiculum 3 days after injecting Nop-tTA mice with TRE-TeTX-YFP virus and continued TTX until harvest at 10 days after viral injection (10 dpi). Although we would have liked to test extended time points in the degenerative process initiated by TeTX, TTX causes locomotor impairment as it diffuses beyond the target region, which limited these experiments to 7 days of infusion. Unlike the GlyCl model, TTX infusion had a marked impact on neuronal morphology in the TeTX mice. Similar to our previous experiments, TeTX mice infused with saline showed pronounced axonal disintegration visible as bright YFP puncta in the DG at 10 dpi (Figure 6A). In contrast, axonal labeling remained robust and intact in mice infused with TTX. Preservation of axonal innervation was limited to the infused hemisphere; degeneration on the contralateral side was identical to that seen in animals infused with saline (Figure 6B). These differences were reflected in both the width of YFP labeling and the integrated intensity of YFP fluorescence measured at the crest of the DG (Figure 6C). Consistent with the axon preservation, the extent of microglial activation normally observed in the DG of TeTX mice was also abated by TTX infusion (Figures 6B-C). The main effects of TTX infusion were localized to the DG, where axonal disintegration precedes cell death markers in the EC. Fractin activation in the EC would just have begun at 10 dpi when these mice were euthanized, and the number of fractin+ cells in the EC was still quite low even in saline-infused mice with no significant differences between conditions (Supplemental Figure 4). If anything, the extent of fractin labeling in neurites – as opposed to soma – appeared qualitatively greater in TTX-infused mice compared to saline, raising the possibility of caspase-assisted neurite remodeling rather than overt cell death (Mukherjee and Williams, 2017; Hollville and Deshmukh, 2018). In sum, TTX infusion preserved axonal innervation and maintained microglial homeostasis in mice expressing TeTX, suggesting that cell death due to synaptic silencing – as opposed to electrical suppression – may be governed by ongoing competition between active and inactive contacts.

**Figure 6.**
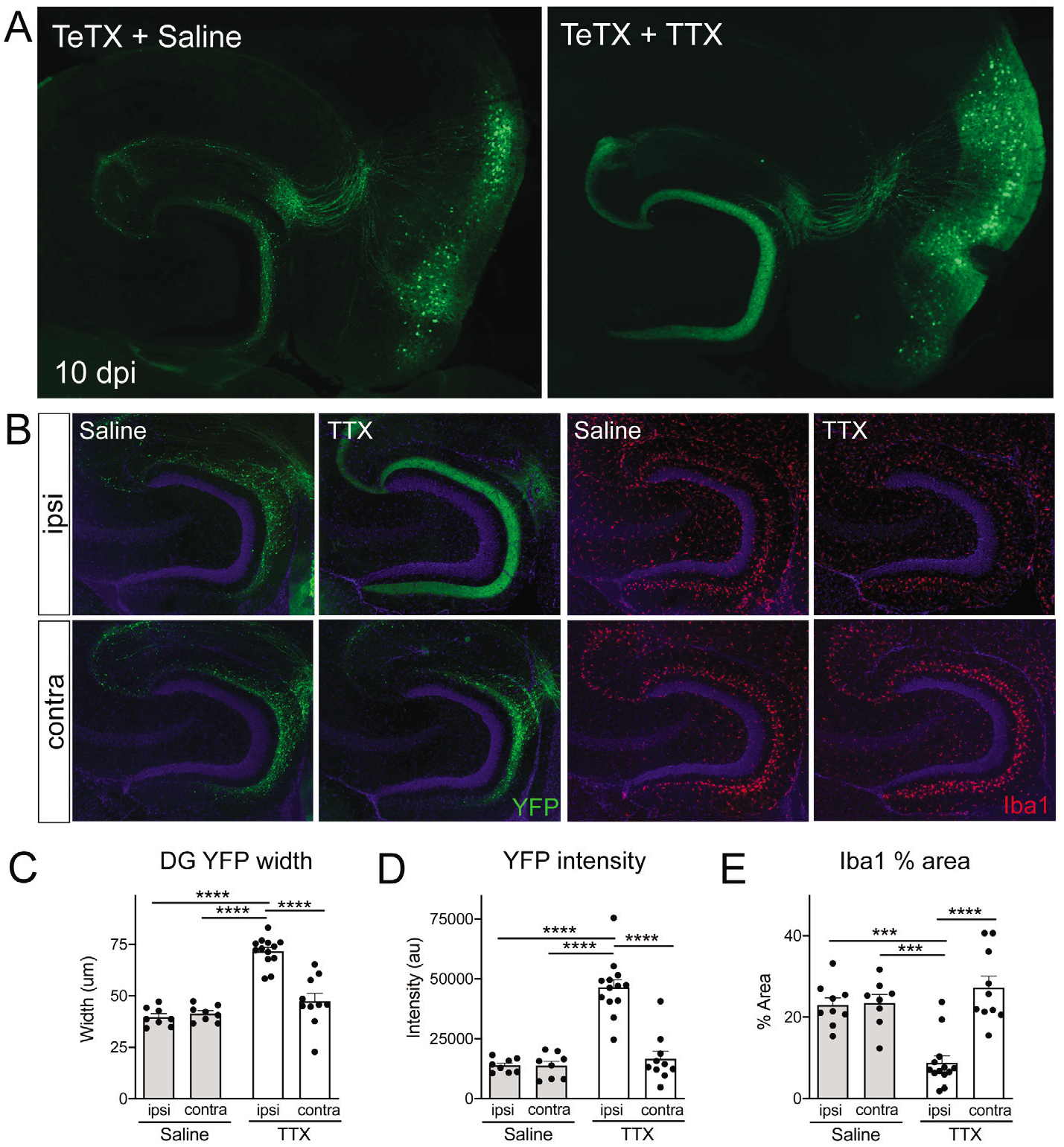
Activity-dependent competition governs EC axonal disintegration due to TeTX silencing. Mice expressing TeTX+YFP in EC neurons were treated by local infusion of TTX or saline. Unilateral infusion targeting the neighboring subiculum began 3 days after viral injection and continued until mice were harvested 7 days later (10 dpi). A. Representative images showing the spread of TeTX transduction in the EC as visualized by native YFP fluorescence. Preservation of DG innervation can be seen in the TTX-infused animal (right), in contrast to the punctate ‘beads on a string’ labeling of remaining perforant path axons in the saline-infused condition (left). B. Left panels: Images show YFP labeling of TeTX-expressing axons (green) in the DG both ipsilateral and contralateral to the infusion cannula. DG innervation is only preserved ipsilateral to TTX infusion. Right panels: Images show Iba1 immunostaining in the DG, where TeTX-evoked microglial activation is prevented in the TTX-infused hemisphere. C, D. The width (C) and intensity (D) of YFP labeling at the crest of the DG confirms axonal sparing ipsilateral to TTX infusion. E. Analysis of % area occupied by Iba1 in the DG confirms that microglial activation was prevented in the TTX-infused hemisphere. Graph data expressed as average values for each animal ± SEM, one-way ANOVA with Tukey’s post-hoc test (missed injection target for some hemispheres precluded 2-way ANOVA). n=8-9 (Saline infusion), 10-13 (TTX infusion). *p<0.05, ***p<0.001, ****p<0.0001.

## Discussion

Our efforts to model entorhinal dysfunction due to Alzheimer’s disease led us to three novel discoveries about this circuit. First, we demonstrate that mature EC2 neurons require ongoing activity for survival long beyond the postnatal period. Second, we demonstrate that electrical vs synaptic silencing can elicit distinct forms of axonal damage prior to cell death. Finally, we show that axonal disintegration due to synaptic silencing is governed by competition between active and inactive neurons, suggesting that this circuit may maintain an activity-dependent critical period well into adulthood. We discuss each of these ideas in turn.

### Activity-dependent neuronal survival persists in the adult cortex

We show that both electrical silencing due to chloride or potassium flux and synaptic silencing due to arrest of neurotransmitter release are each sufficient to cause axonal damage of EC2 neurons followed by cell body demise. Such wholesale changes in activity-dependent circuit patterning are traditionally thought to be limited to a brief window during postnatal development (Murase, 2014; Wong and Marin, 2019). Because our experiments focused primarily on the EC, we do not know the extent to which activity-dependent neuronal survival exists outside of this circuit; however, past work has uncovered both experimental manipulations and traumatic conditions that are capable of inducing activity-dependent axonal remodeling in the adult cortex (Fox and Wong, 2005; Dancause, 2006; Spolidoro et al., 2009; Bavelier et al., 2010). We suspect that our discovery arises at least in part from three fortuitous aspects of our experimental system: 1) the prolonged suppression provided by the manipulations we chose compared with the shorter duration tools used in earlier work, 2) the decision to look at time points well after silencing had subsided, and 3) our focus on the EC-DG connection where circuit architecture, a layer-specific driver line, and a co-expressed cytosolic YFP label allowed clear visualization of axonal degeneration and cell loss.

### Prolonged silencing uncovered activity-dependence

We were fortunate to study EC2 as it allowed us to clearly identify perforant path axons, their terminals in DG, and their cell bodies in the cortex. Other cortical circuits lacking this discreet organization would not have provided such a clear view of circuit damage. Nevertheless, numerous studies have used chemical, chemogenetic, or optogenetic tools to silence entorhinal input to the DG without observing signs of axonal damage or cell death (Miao et al., 2015; Ormond and McNaughton, 2015; Rueckemann et al., 2016; Kanter et al., 2017; Robinson et al., 2017; Qin et al., 2018; Rowland et al., 2018; Heys et al., 2020; Rodriguez et al., 2020; van Wijngaarden et al., 2020). These studies have targeted EC2 with varying degrees of specificity, from broad chemical inactivation via muscimol infusion or hSyn-controlled AAV transduction to more targeted silencing using the same Nop-tTA driver studied here (i.e., EC-tTA). Nearly all of these studies used EC inactivation to test the electrophysiological consequences of entorhinal dysfunction on spatial coding and spatial navigation behaviors. This focus lead to two key differences that may explain why activity-dependent neuronal survival was not observed in these previous studies. First, changes in the desired outcomes, such as altered place cell properties, can be detected immediately after entorhinal inactivation, and animals were generally harvested upon completion of the task. We delayed harvesting the initial Nop-tTA; TRE-GlyCl-YFP mice until many days after IVM injection to map the pharmacodynamics of extended silencing with the GlyCl system. Second, optogenetic or DREADD chemogenetic tools display temporal precision on the scale of milliseconds to minutes, but both GlyCl and TeTX systems for neuronal silencing are slow and sustained. Although the effects of GlyCl, DREADD, and optogenetic silencing are all transient, the half-life for IVM in the brain is over 13 hours and behavioral impairments persist for several days after IVM administration ((Lerchner et al., 2007; Zhao et al., 2016) and data not shown). The effect of TeTX is even more protracted, in theory silencing transmission as long as expression continues. While neither the GlyCl or TeTX systems would be ideal for experiments requiring restoration of physiology or behavior after silencing, both were well suited to our initial purpose of modeling circuit impairments due to neurodegenerative disease. The prolonged neuronal silencing achieved by GlyCl and TeTX was critical to our observations, and if the pharmacodynamic studies had not included timepoints many days after the initial IVM injection, we would have missed the dramatic changes in YFP+ labeling.

One previous experiment examined chronic chemogenetic EC inactivation rather than acute silencing. Rodriguez et al. used viral expression of hM4Di DREADD combined with chronic CNO infusion via osmotic pump to persistently suppress firing in EC neurons over six weeks of study (Rodriguez et al., 2020). This paper is most similar to our experiments in its prolonged manipulation of EC activity, but focused on downstream tau and amyloid pathology in the hippocampus rather than cell survival or axonal morphology in the EC. Immunostaining for virally-expressing cells following 6 weeks of CNO infusion suggested that many DREADD-expressing neurons survived. However, we would be fascinated to know how the numbers of surviving neurons compared to the number of DREADD-positive EC2 cells present before the prolonged silencing.

### Different modes of silencing elicited distinct patterns of axonal damage

We unexpectedly discovered that the same population of EC2 neurons could adopt different patterns of axonal degeneration prior to cell death. Axons silenced by GlyCl displayed evidence of simple axonal retraction beginning within 1-2 days of silencing onset. These axons remained intact, although displaced from the DG, with bulb-like structures at their tips. This pattern was strikingly different from the overt axonal disintegration observed after TeTX silencing. Atrophy in TeTX-expressing axons was reminiscent of Wallerian degeneration, which is characterized by axon blebbing and fragmentation. Both patterns of axon elimination are found in the developing CNS, although rarely in the same cell population (Neukomm and Freeman, 2014; Riccomagno and Kolodkin, 2015). The developing thalamocortical projection stands out in displaying both retraction and disintegration, with the choice seeming to depend on the length of axon segment being remodeled (Portera-Cailliau et al., 2005; Neukomm and Freeman, 2014). Axon length did not differ in our model, as the same projections underwent degeneration under each form of silencing. Instead, our models differed by the method used to prevent neurotransmission. GlyCl suppresses electrical activity by membrane hyperpolarization (Lerchner et al., 2007; Zhao et al., 2016) or reduced membrane conductance (Weir et al., 2017), either of which would dampen but not eliminate evoked neurotransmitter release onto postsynaptic DG targets while having no effect on spontaneous release. In contrast, TeTX has no direct effect on electrical activity, but can potently inhibit both evoked and spontaneous neurotransmitter release by preventing vesicle fusion at the membrane (Schiavo et al., 1992; Humeau et al., 2000). Future studies will need to address the distinct mechanisms driving the axonal degeneration observed by these two different means of neuronal silencing.

### Neuronal competition regulates axon degeneration due to TeTX but not cell loss due to GlyCl

Competition between axons determines which cells live or die during neural development (Blakemore, 1991; Zhao and Reed, 2001; Yu et al., 2004). While competitive processes have long been studied in central sensory systems, only recently was this mechanism found to govern axonal refinement in the developing perforant pathway (Yasuda et al., 2011). This study revealed that EC2 axons silenced with TeTX were eliminated from the circuit within a week of reaching the DG. They next found that infusing TTX into the EC of TeTX-expressing mice - to prevent all activity through the perforant path - prevented degeneration. This elegant experiment suggested that competition between active and inactive axons drove developmental refinement of the perforant path. We used a similar approach to determine if competition might also drive axonal degeneration and cell death in the adult EC. The divergent outcome of this manipulation in GlyCl and TeTX models surprised us, but supported the idea that synaptic silencing and electrical suppression evoke distinct mechanisms of cell death in the adult EC. Local infusion of TTX preserved axonal integrity in neurons expressing TeTX, but had no effect on cell loss in neurons silenced by GlyCl. Electrical silencing by GlyCl appears to kill EC2 neurons by a non-competitive, possibly cell-intrinsic mechanism, but axonal disintegration due to synaptic silencing by TeTX appears to be influenced by activity in neighboring neurons. This competitive mechanism – and the experiments used to demonstrate it - are consistent with the process used during developmental refinement of the perforant path; however, we cannot can rule out the alternative explanation that TTX may act directly on the TeTX-expressing cells rather than through their neighbors. Our experiments also stop short of demonstrating that axonal preservation in TeTX-expressing mice is sufficient to protect the cell body, due to limitations on the duration of TTX infusion. Nevertheless, this discovery raises the possibility that the critical window for EC circuit refinement - potentially including wholesale neuron elimination - remains open long into adulthood.

That synaptic silencing can evoke cell death and uncover the possibility of latent competition in the perforant path raises the opposing question of whether increasing synaptic activity might have a protective effect. More precisely, it is possible that sparsely increasing neurotransmitter release in the EC could protect the more active cells, but induce cell death in their neighbors. During brain development, increasing activity through environmental or pharmacologic stimulation causes enlargement of the stimulated axonal field and increased survival of the stimulated neurons (Assali et al., 2014; Blanquie et al., 2017). Several groups have tested EC activation in the adult brain using chemo- or optogenetic tools. However, most studies used these tools to evoke short bursts of activity that may not be sufficient to induce a survival imbalance (Zhang et al., 2013; Zhang et al., 2014; Kanter et al., 2017; Leung et al., 2018). Two studies performed repeated daily EC activation via DREADDs or channelrhodopsin, but tested behavioral and histopathological outcomes rather than neuroanatomical features (Yamamoto et al., 2015; Yun et al., 2018). Unfortunately, neither had reason to examine how their EC manipulation affected the integrity of non-activated neighboring cells.

### Limitations of our experimental models

We also must address limitations of our model systems which introduce important caveats to several of our conclusions. First and foremost, our studies focused almost exclusively on EC2. We do not know if our chemogenetic tools would evoke axonal damage and cell death in other areas of the brain. We tried to address this concern by measuring cell loss in neural populations outside EC that were also silenced in the Nop-GlyCl model. Although we found no evidence of cell loss in presubiculum, parasubiculum, or retrosplenial cortex, we also acknowledge that these regions likely expressed lower levels of GlyCl than EC2 in our model. Both the intensity of YFP fluorescence and the efficiency of IVM-evoked firing suppression were greatest in EC2, followed by parasubiculum, presubiculum, and retrosplenial cortex. We note that fractin staining and microglial activation were occasionally found in border between parasubiculum and EC and that doubling the dose of IVM caused both markers to increase in a subset of mice (data not shown), although we did not observe significant cell loss in these neighboring regions. Nevertheless, this finding suggests that EC2 may not be alone in sensitivity to chloride-based silencing and that other cell types may become vulnerable at higher doses or with greater GlyCl expression.

We also appreciate that our supporting evidence for EC2 cell death provided by viral expression of Kir, hM4Di, and TeTX was subject to variability of viral delivery between animals that prevented us from performing meaningful cell counts at increasing dpi. These experiments instead relied on loss of DG innervation and the appearance of damage markers such as Iba1 and fractin to support our conclusions; however, these outcomes are not as definitive as the decreased cell number observed in the GlyCl system. This concern is offset in the case of TeTX by the physical axon disintegration which develops alongside these surrogate markers. However, this observation contradicts past work where TeTX was used for silencing other pathways of the trisynaptic loop for up to 6 months with no evidence of lasting axonal damage (Nakashiba et al., 2008; Lopez et al., 2012; Nakashiba et al., 2012). Of course, our experiments placed TeTX in EC2 neurons, while the Tonegawa studies expressed TeTX in DG and CA3 neurons. We also expressed TeTX using a viral construct, while the DICE system relied on germline transgenic constructs. We do not know whether these distinctions are relevant, but we also note that EC2 appears vulnerable to a variety of viral and transgenic manipulations well beyond TeTX.

### Vulnerability of the EC-hippocampal circuit may derive from its function

While we cannot rule out the possibility that prolonged silencing may cause cell death in other areas of the brain, EC2 stands out in several respects which drove our original interest in studying this pathway. Unlike the primary sensory cortices, which precisely map their input during a defined critical period, the EC remains highly plastic throughout life and is constantly modified during memory formation. This need to remain continuously plastic may render the EC more vulnerable to insults that cause an activity imbalance between neurons. Here we used several methods to artificially impair transmission between the EC and hippocampus, but several naturally-occurring insults may have similar consequences. For example, formation of neurofibrillary tangles in EC neurons might impair the initiation or propagation of action potentials in affected cells while neighboring neurons remain normally active. EC2 is particularly prone to tangle pathology, where these lesions mark the earliest stages of Alzheimer’s disease (Hyman et al., 1984; Van Hoesen et al., 1986; Braak and Braak, 1991; Van Hoesen et al., 1991; Braak and Braak, 1995). Alternatively, the formation of amyloid plaques in the DG could physically disrupt the communication between pre- and post-synaptic cells. Plaques are prominent in the molecular layer of the dentate, precisely where the presynaptic terminals of the perforant path connect with postsynaptic dendrites of granule cells (Geddes et al., 1986; Hyman et al., 1986; Crain and Burger, 1988; Braak and Braak, 1991). Importantly, in the early stages of pathology formation, these communication deficits would be localized to individual neurons with tangles or their axons terminating near plaques. The altered microenvironment of these affected neurons could establish a competitive situation in which those neurons would be disadvantaged relative to their neighbors, perhaps leading to their demise. This theory offers a new model by which the pathologies of Alzheimer’s disease may underlie the early vulnerability of EC neurons. Moreover, these studies raise the possibility that lifelong plasticity needed for episodic memory may be a disadvantage in disease.

## Methods

### Mice

Animals were housed in accordance with the NIH Guide for the Care and Use of Laboratory Animals, and protocols approved by the Baylor College of Medicine Institutional Animal Use and Care Committee. All experimental animals were three- to six-months of age at the time of treatment. Mice of both sexes were used for all experiments.

Nop-tTA Line S (also known as tTA-EC Prss19 (Yasuda and Mayford, 2006), MMRRC strain #31779) and TetO-GFP-nls-LacZ (also known as nls-lac-CMK (Mayford et al., 1996)) were gifts from Mark Mayford, Scripps Research Institute. The lines were maintained on a C57BL/6J background.

TRE-GlyCl-YFP Line 9531 ((Zhao et al., 2016) Jax strain #29301) was described previously and was maintained on a FVB/NJ background for more than 10 generations.

TRE-hM4Di mice were obtained from Jackson Laboratory, stock # 024114 (Zhu et al., 2014) and maintained on a C57BL/6J background.

Nop-GlyCl and their single transgenic Nop-tTA siblings for these experiments were derived from the intercross of male Nop-tTA x female TRE-GlyCl breeders (F1 FVB;B6 background). Similarly, Nop-hM4Di animals were derived from intercross of Nop-tTA x TRE-hM4Di animals (C57BL/6J background).

Nop-tTA mice used for viral injections were derived from the intercross of male Nop-tTA x female FVB/NJ wildtype breeders (Jax #1800) or from the single transgenic offspring of male Nop-tTA x female TRE-GlyCl breeders (both resulting in an F1 FVB;B6 background).

### Viral constructs

Four tTA-dependent pAAV expression vectors were created for study, one encoding YFP alone, one encoding tetanus toxin light chain (TeTX) with YFP, one encoding Kir with YFP, and one encoding hM4Di with YFP. pAAV-TRE3G-T2a-YFP contained the TRE3G promoter from pTRE3G (#631168, Clontech, Mountain View, CA), followed in turn by a modified multiple cloning site (BamHI-MfeI-AgeI-NruI-SpeI), the *Thosea asigna* virus 2A sequence, the enhanced yellow fluorescent protein (YFP) from pEYFP-N1 (#6006-1, Clontech), the woodchuck hepatitis virus posttranscriptional regulatory element (WPRE, from pAAV2-GluClα-WPRE (Lerchner et al., 2007)), and a bovine growth hormone (BGH) polyadenylation signal. The entire insert was flanked by inverted terminal repeats (ITR) from AAV2.

Tetanus toxin light chain (TeTxLC) was amplified by PCR from pGEMTEZ-TeTxLC (Addgene #32640, (Yu et al., 2004)) using forward primer: GCGCGCAATTGGCCACCATGCCGATCACCATCAACAACTT and reverse primer: GCGCGACTAGTAGCGGTACGGTTGTACAGGTT. The resulting PCR product was digested with MfeI and SpeI and ligated into the MfeI and SpeI sites of pAAV-TRE3G-T2a-YFP to build pAAV-TRE3G-YFP-T2a-TeTxLC.

Kir was amplified by PCR from pCAG-Myc-Kir2.1^E224G/Y242F^-2A-EGFP (similar to pCAG-Kir2.1-T2A-tdTomato, Addgene #60598, kind gift of Mingshan Xue, (Xue et al., 2014)) using forward primer: GCGCGGGATCCGCCACCATGGAGCAGAAGCT and reverse primer: GCGCGACTAGTTCCACTGCCTATCTCCGATTC. The resulting PCR product was digested with BamHI and SpeI and ligated into the BamHI and SpeI sites of pAAV-TRE3G-T2a-YFP to build pAAV-TRE3G-YFP-T2a-Kir^E224G/Y242F^.

hM4D(Gi) was amplified by PCR from pcDNA5/FRT-HA-hM4D(Gi) (Addgene #45548 (Armbruster et al., 2007)) using forward primer: GCGCGGGATCCGCCACCATGGCCAACTTCACACCTGTC and reverse primer: GCGCGACTAGTCCTGGCAGTGCCGATGTTC. The resulting hM4D(Gi) fragment was digested with MfeI and SpeI and ligated into pAAV-TRE3G-NMCS-YFP (plasmid available from the Jankowsky lab) to create pAAV-TRE3G-hM4Di-T2a-hM4Di.

### Viral packaging

Packaging and purification of viral constructs in AAV8 was performed by the Gene Vector Core at Baylor College of Medicine, as described previously (Huichalaf et al., 2019; Park et al., 2021)

### IVM administration

GlyCl-expressing bigenic or trigenic mice and controls were administered with either vehicle solution (60:40 propylene glycol: isotonic saline) or 5 mg/kg ivermectin (1% Ivomec, Merial, Lyon, France, or 1% Noromectin, Norbrook, Newry, Northern Ireland) diluted to 2 mg/ml in vehicle) via intraperitoneal injection. Mice were scarified for histological assessment at time points ranging from 6 hr to 30 d after treatment, as indicated. For 2x ivermectin treatment, GlyCl expressing animals received 10 mg/kg ivermectin (or comparable volume of vehicle) at 2 mg/ml.

### Stereotaxic viral delivery

Adult Nop-tTA mice on a F1 FVBB6 background underwent surgery between 3-5 months of age. Mice were fixed in a stereotaxic frame, administered buprenorphine, ketoprofen, and local lidocaine/bupivacaine for analgesia, and anesthetized with isoflurane. Bilateral craniotomies were made using a 0.45 mm drill bit. Craniotomies targeting the EC were positioned at ML ±3.60 mm, while AP coordinates were adjusted for each animal using the bregma-lambda distance (right hemisphere: AP = 0.808 × BL distance + 1.636 mm; left hemisphere: AP = 0.829 × BL distance + 1.46 mm). For EC expression, AAV8-TRE3G-YFP, AAV8-TRE3G-TeTX_LC_-YFP, or AAV8-TRE3G-Kir2.1-YFP was injected bilaterally using a 31 gauge stainless steel needle (#22031-01, Hamilton Co., Reno, NV) attached to a microsyringe injector (#UMC4, World Precision Instruments, Sarasota, FL). The needle was slowly lowered to DV −3.2 mm and remained in place for 10 min after the injection. Each hemisphere was injected with 300 nl virus (6.9 × 10^8^ genome copies total for YFP and TeTX, empirically equivalent to 5.7 × 10^8^ gc for Kir2.1) at a rate of 50 nl/min.

### P0 viral injection

Intracerebral viral injection was performed as previously described (Kim et al., 2014; Kim et al., 2016; Huichalaf et al., 2019). Neonatal Nop-tTA animals were stereotaxically injected at postnatal day 0 with 1 ul/hemisphere of AAV8 at a rate1 ul/min (5.4 × 10^9^ gc/ul of TRE3G-hM4Di-T2a-YFP), targeting the lateral ventricles bilaterally: AP −1.0; ML +/- 1.35; DV −1.7.

### CNO administration

Clozapine N-oxide (CNO; ENZO, #BML-NS105) was dissolved in sterile 0.9% saline at a concentration of 8 mg/ml. The mini-osmotic pump (Alzet, #2001D, flow rate 8 ul/hr) was filled with 200 ul of CNO solution or saline and primed in sterile saline for 2 hr at 37 °C. The primed mini-osmotic pump was inserted subcutaneously along the flank of anesthetized mice to deliver 1.5 mg CNO/day (approximately 50 mg/kg/d). Pumps were replaced every 24 hr for two d (3 pumps total). The mice were euthanized and brains dissected 7 d after initial CNO exposure.

### Optimizing TTX dosage, targeting, and spread

TTX (Abcam, Cambridge, MA, ab120054) or 0.9% saline containing 0.04% Trypan blue for visual confirmation of catheter flow was pre-loaded into a micro-osmotic pump (#1003D, Alzet, Cupertino, CA) and primed by immersion in sterile 0.9% saline overnight at 37 °C. Mice were anesthetized with isoflurane, and an indwelling 30 gauge cannula (#0008851, Alzet) cemented to the skull (Kerr, Orange, CA, 34417) at AP −4.5, ML +3.0 and DV −2.5 mm to target immediately above the right subiculum or at AP −3.1, ML +3.0, DV −2.65 to target the dorsal DG. Immediately following cannula placement, the primed micro-osmotic pump was positioned subcutaneously through a mid-scapular incision and the catheter routed subcutaneously to connect with the cannula. The pump flow rate of 1 μl/hr was chosen to deliver 24 μl/day TTX. Mice were injected with pentylenetetrazol (PTZ, 25 mg/kg) 90 min prior to harvest and killed at 6-48 hr after implantation to examine the spread of dye and the extent of c-fos labeling induced by PTZ.

### TTX administration

TTX (2.3 uM) or 0.9% saline containing 0.04% Trypan blue was prepared and infused as described above to target above the right subiculum. Where indicated, 12-24 hr post-surgery, mice received a single i.p. injection of IVM (5 mg/kg) or vehicle. For Nop-GlyCl mice, the osmotic pump was removed 3 d after implantation and animals were harvested 7 d after IVM or vehicle injection. For TeTX mice, the osmotic pump was implanted 3 d after viral injection and replaced 4 and 7 d later to continue TTX administration until animals were harvested 10 d after viral injection.

### Tissue harvest and sectioning

Mice were killed by pentobarbital overdose and transcardially perfused with phosphate buffered saline (PBS) followed by 4% paraformaldehyde (PFA) in PBS. The brain was removed and post-fixed by immersion in PBS containing 4% PFA at 4 °C overnight, and then cryoprotected by immersion in 30% sucrose in PBS at 4 °C until equilibrated. Tissue was sectioned in the horizontal plane (for RSC, pre/parasubiculum, and entorhinal staining) or the coronal plane (for substantia nigra staining) at 35 μm thickness using a freezing-sliding microtome. Sections were stored in cryoprotectant media at −20 °C until use.

### Immunolabeling and histology

#### Fractin and YFP, fluorescence

For labeling of fractin with or without co-detection of YFP, free-floating sections were rinsed in TBS and then endogenous peroxide was quenched by incubating sections 30 min at RT with 0.9% H_2_O_2_ in TBS with 0.3% Triton X-100 and 0.05% Tween 20. Sections were washed in TBS, blocked with 5% normal goat serum in TBS with 0.3% Triton X-100 and 0.05% Tween 20 for 1 hr at RT, and then incubated in primary antibody at 4 °C for 72 hr (1:1,000 chicken anti-GFP, Abcam, ab13970; 1:5000 rabbit anti-fractin, Phosphosolutions, 592-FRAC). Sections were washed several times in TBS, followed by incubation overnight at 4 °C in secondary antibody diluted in block (1:500 goat anti-chicken Alexa 488, Life Technologies, A11039; 1:5000 biotinylated goat anti-rabbit, Vectastain Elite ABC HRP kit, PK-6101, Vector Laboratories, Burlingame, CA). Sections were washed several times in TBS with 0.3% Triton X-100 and 0.05% Tween 20, incubated for 30 min at RT with Vectastain components A and B diluted 1:30 in TBS, and then washed overnight in TBS. Sections were incubated for 10 min in Cy3 Plus Amplification Reagent diluted 1:300 in 1x Plus Amplification Diluent (TSA Plus Cyanine 3 System, Perkin Elmer, Waltham, MA, NEL744001KT), washed several times in TBS with 0.3% Triton X-100 and 0.05% Tween 20, incubated 10 min in 0.2 ug/ml DAPI diluted in TBS, and then washed again in TBS.

#### Cleaved caspase 3 and YFP, fluorescence

Sections were rinsed with TBS and blocked with TBS containing 0.1% Triton X-100 (TBST) plus 5% normal goat serum for 1 hr at RT before 3 d incubation in primary antibody diluted in block (1:1,000, chicken anti-GFP, Abcam, ab13970; 1:200 rabbit anti-cleaved caspase 3, EMD Millipore, AB3623). Sections were washed several times in TBS, followed by overnight incubation at 4 °C in secondary antibody diluted 1:500 in block (donkey anti-rabbit Alexa 568, Life Technologies A10042). Sections were again washed in TBS. Tissues were then incubated overnight at 4 °C in tertiary antibody diluted in block (goat anti-chicken Alexa 488, Life Technologies, A11039, and goat anti-donkey Alexa 568, Life Technologies (discontinued)), and washed a final time in TBS.

#### Iba1 and YFP, fluorescence

For fluorescence labeling of Iba1 with or without co-detection of YFP, sections were rinsed with TBS and then blocked with TBST plus 5% normal goat serum for 1 hr at RT before overnight incubation in primary antibody diluted 1:1000 in block (rabbit anti-Iba1, Wako Chemicals USA, Richmond, VA, 019-19741; 1:1,000 chicken anti-GFP, Abcam). Sections were washed several times in TBS, followed by 2 hr incubation at RT in secondary antibody diluted 1:500 in block (goat anti-rabbit Alexa 568, Life Technologies; 1:500 goat anti-chicken Alexa 488, Life Technologies). Sections were again washed in TBS, incubated 10 min in 0.2 ug/ml DAPI diluted in TBS before being mounted and coverslipped.

All sections were mounted onto Superfrost Plus slides (Fisher Scientific, Pittsburgh, PA) and coverslipped with ProLong Diamond anti-fade mounting media (P36970, Invitrogen/Life Technologies Corp, Eugene, OR).

#### Iba1, colorimetric

Sections were rinsed with TBS before endogenous peroxide was quenched by incubating sections for 30 min at RT in TBS with 0.9% hydrogen peroxide and 0.01% Triton-X-100. Sections were washed in TBS and then blocked with TBS containing 0.1% Triton X-100 (TBST) plus 5% normal goat serum for 1 hr at RT before overnight incubation in primary antibody diluted 1:1000 in block (rabbit anti-Iba1, Wako Chemicals USA). Sections were washed several times in TBS, followed by overnight incubation at 4 °C in biotinylated secondary antibody diluted 1:500 in block (biotinylated goat anti-rabbit, Vectastain Elite ABC HRP kit, PK-6101, Vector Laboratories). Sections were washed in TBS and then incubated for 1 hour at RT in Vectastain kit components A and B, each diluted 1:10 in TBS. Sections were again washed and then developed for 1 hour in DAB solution (Sigmafast D4418 tablets dissolved in 50 ml water). Sections were washed in TBS, mounted on Superfrost Plus slides, and allowed to dry overnight. Slides were processed through alcohol series (70%, 95%, 100%) to xylene and then coverslipped with Permount (Fisher Scientific, Pittsburgh, PA).

#### β-galactosidase stain

Bigenic Nop-tTA/tetO-LacZ and trigenic Nop-tTA/tetO-LacZ/TRE-GlyCl tissues were harvested and sectioned as above, except that brains were post-fixed by PFA immersion for 2 hr prior to immersion in 30% sucrose in PBS. A 1 in 6 series of sections was washed in TBS, mounted onto Superfrost Plus slides, and allowed to dry for 24-72 hr. Slides were rehydrated in solution A (2 mM MgCl_2_ in PBS) for 35 min, incubated in pre-warmed solution B (2 mM MgCl_2_, 0.2% NP40, 0.1% sodium deoxycholate in PBS) for 1 hour, and then incubated in pre-warmed solution C overnight (solution B with 10 mM ferro/ferri cyanide, 0.6 mg/mL X-gal). Slides were rinsed in stop solution (0.1% Triton-X 100, 1 mM EDTA in HEPES buffered saline, pH 7), post-fixed overnight with 4% PFA, washed in PBS, and then air dried for 1 hr. Slides were dehydrated through alcohol series (70, 95%, 100%) to xylene, followed by rehydration to water. Slides were counterstained in nuclear fast red, dehydrated again through the alcohol series to xylene, and finally coverslipped with Permount.

### Imaging and image quantification

All figure images have been taken at exposure times that optimize the display histogram and then adjusted in Photoshop to visually match the background levels across animals. All images used for quantitation within each experiment were imaged at the same exposure and adjusted identically for analysis.

#### Fractin, YFP+ cells, YFP+ retraction bulbs

A 1 in 6 series of horizontal sections through the entire dorso-ventral extent of the brain were co-immunostained for YFP and fractin as described above. Tiled images were acquired using a Zeiss Axio Scan.Z1 at 10x magnification (Carl Zeiss AG, Oberkochen, Germany). Exposure time and lamp intensity were constant for all sections. Sections used for analysis were plane-matched across animals. Regions of interest were identified from anatomical landmarks, and objects within each ROI were manually counted using the multi-point tool and ROI manager in ImageJ/Fiji Mac v2.0.0. For Nop-GlyCl animals without TTX treatment (Figure 1), YFP+ and fractin+ cells were counted in EC layer 2 from four horizontal sections spaced at ~300 um intervals spanning −2.68 to −3.60 mm from bregma, (Franklin and Paxinos, 2008). For Nop-GlyCl and Nop-tTA+TeTX animals with TTX treatment (Figures 5, 6, and Supplemental Figure 4) six to eight sections spanning −2.68 to −3.60 mm from bregma were used for analysis and all fractin+ cells were manually counted using both hemispheres for analysis. Retraction bulbs were counted in the deep layers of EC (below layer 2) in four horizontal sections across the same interval (Figure 1). Graph data expressed as average values per section for each animal ± SEM

#### β-galactosidase

A 1 in 6 series of horizontal sections through the entire dorso-ventral extent of the brain was stained for lacZ as described above. Tiled images were acquired using a Zeiss Axio Scan.Z1. Regions of interest were identified using anatomical landmarks and blue-labeled nuclei within the ROI were manually counted using the multipoint tool and ROI manager in ImageJ/Fiji Mac v2.0.0. For the EC, labeled nuclei were counted bilaterally in all sections spanning the entire EC2 (6-8 sections per mouse). For the pre- and parasubiculum, lacZ+ nuclei were counted bilaterally for three sections spanning −2.36 to −2.68 mm along the dorso-ventral axis. Graph data expressed as the total bilateral cell count from each animal ± SEM. For the RSC, we initially counted labeled nuclei within a single section at −1.24 mm from bregma for all animals. This value was used to validate automated analysis of percent area covered by lacZ staining in the same section after binary thresholding using the Otsu method. Upon confirming the equivalence of manual cell counting to automated area analysis, three sections starting at −1.24 from bregma were analyzed using the automated assessment of percent area stained by lacZ. Graph data expressed as average % area for each animal ± SEM.

#### Iba1 and YFP % area

A 1 in 6 series horizontal sections stained for Iba1 was scanned using a Zeiss Axioscan Z.1 and imported into Image J2 ((Fiji Is Just) ImageJ 2.0.0-rc-68/1.52g; Java 1.8.0_172 (64 bit)). From these, three (fluorescence (Figures 4 and 5)) or four (histochemistry, Figure 1) plane-matched sections spanning the region of −2.36 to −4.12 mm from bregma and 3.44 to 1.68 mm interaural (plates 144-155 in Franklin and Paxinos, 3^rd^ edition (Franklin and Paxinos, 2008)) were chosen for analysis (three plane-matched sections were used for dpi 10-13). Each image was converted to 8-bit and three regions of interest were manually outlined: superficial EC covering layers 2-3, molecular layer of the DG including both blades, and the medial zone of stratum radiatum in CA1. CA1 was used to determine the background threshold for each section one hemisphere at a time using the Yen method. The percent area above the CA1 background was then measured in EC and DG. Values from both hemispheres were averaged for Iba1 area shown in Figure 1 and Supplemental Figure 3, or analyzed separately for data shown in Figures 5 and 6.

#### YFP intensity, DG width

For the TeTX+TTX experiment shown in Figure 6, the width of YFP labeling was measured across the crest of the dentate gyrus using the measure feature in Fiji. YFP immunofluorescence intensity was measured using the integrated density function of Fiji from a 50 μm x 50 μm region within the crest of the DG (IntDen = area x mean gray value). Regional values were measured from each hemisphere individually in three plane-matched sections per animal and averaged to calculate YFP intensity. For the Kir experiment shown in Supplemental Figure 2, YFP immunolabeling within the DG was manually outlined and measured using the mean gray value of Fiji (MGV = sum of pixel values / number of pixels). This measure accounts for differences in the size of selected ROIs. Regional values were measured from two plane-matched sections per animal and averaged to calculate YFP intensity.

### Brain slice preparation and electrophysiology

Brain slices from Nop-GlyCl or virally-injected Nop-tTA mice (YFP, TeTX-YFP, or Kir-YFP) were prepared as described previously with minor modification (Li et al., 2015; Zhao et al., 2016; Li et al., 2017). Mice were deeply anesthetized with isoflurane and the brains were rapidly removed and placed in N-methyl-D-glucamine (NMDG) based “cutting buffer” containing (in mM): 80 NMDG, 2.5 KCl, 0.5 CaCl_2_, 10 MgSO_4_, 1.2 NaH_2_PO_4_, 30 NaHCO_3_, 25 D-Dextrose, 5 Sodium ascorbate, 2 Thiourea, 3 Sodium pyruvate, pH 7.4 adjusted with HCl and 300-310 mOsm (Ting et al., 2014). Coronal or horizontal slices containing EC, presubiculum, parasubiculum, or retrosplenial cortex (300-320 μm) were cut in 95% O_2_- and 5% CO_2_-oxygenated cutting buffer. Slices were incubated at 35 °C in a submerged chamber containing artificial cerebrospinal fluid (aCSF) equilibrated with 95% O_2_ + 5% CO_2_-oxygenated buffer for at least 30 min and maintained at room temperature afterward until transfer to a recording chamber. The aCSF contained (in mM): 126 NaCl, 2.5 KCl, 2.4 CaCl_2_, 1.2 MgCl_2_, 1.2 NaH_2_PO_4_, 21.4 NaHCO_3_, 11.1 D-Dextrose, pH 7.4, and the osmolarity was adjusted to 300-310 mOsm.

Whole-cell patch-clamp recordings were made from visually identified YFP-labeled or unlabeled EC layer 2, presubiculum, parasubiculum, or retrosplenial neurons with infrared DIC and fluorescence optics (AxioExaminer, Zeiss). Recording electrodes were pulled from borosilicate glass (1.5 mm x 0.86 mm diameter) on a P-1000 horizontal micropipette puller (Sutter Instruments) to a resistance of 4-5 MΩ when filled with the intracellular solution. Current-clamp experiments used an intracellular pipette solution containing (in mM): 120 K gluconate, 20 KCl, 0.1 EGTA, 4 MgCl_2_, 5 NaCl, 10 HEPES, 4 MgATP, 3 NaGTP, 10 phosphocreatine, pH 7.2, 290 mOsm. Data were low-pass filtered at 5 kHz and sampled at 10 kHz using Axon MultiClamp 700 B amplifier and Digidata 1440A data acquisition system under control of pClamp 10.7 software (Molecular Devices). Recordings measured the number of action potentials generated in response to 250 ms stepped current injections (−60 pA to 200 pA at 20 pA increments). Throughout the experiment, access resistance (Ra) was 10 – 20 MΩ and cells were discarded if Ra drifted above 20%. Series resistance and cell capacitance were compensated. Current-response relationships were constructed by counting the number of action potentials generated at each current step in the presence or absence of 100 nM IVM. IVM was allowed to perfuse for 10-15 min before post-IVM recordings were made. All recordings were made in ACSF at an ambient temperature of 32 – 34 °C. One neuron was recorded per slice and four to five slices were recorded per mouse. Results compared pre- and post-IVM for each neuron.

In a separate set of experiments, extracellular field potentials were recorded using electrodes filled with aCSF described above. Excitatory postsynaptic potentials (fEPSPs) were recorded in the DG medial molecular layer following perforant path stimulation through a twisted nichrome wire electrode. Stimulus-response curves were calculated by plotting initial slope of the fEPSPs against stimulus intensity (50-500 uA in 25 uA steps). All recordings were made at ambient temperature of 32 – 34 °C.

### Statistical analysis

Data comparisons were done using two-tailed unpaired Students t-test for two group comparisons, or one-, two-, or three-way ANOVA for three or more group comparisons, followed by Dunnett’s (one-way for comparison to control), Tukey’s (one-way for comparison across all groups), or Sidak’s (two-way RM for comparison to control and three-way for comparison across all groups) post-test. Statistical analyses and graphing of data were done using Prism 6.0 or 8.4 (GraphPad, La Jolla, CA). All graphs display group means ± SEM.

## Acknowledgements

We thank Jennifer Boenker, M. Danish Uddin, Shaina Gong, and Rebecca Corrigan for assistance with animal care and genotyping; Mingshan Xue and Daoyun Ji for valuable experimental advice; and Mark Mayford and Karsten Baumgartel for sharing the Nop-tTA and nls-lac-CMK mouse lines. This work was funded by NIH RF1 AG058188, RF1 AG058188-01S1, RF1 AG054160, and R01 NS092615 to JLJ, R25 GM056929 (support for GAP), F31 AG067676 to CAW, Alzheimer’s Association fellowship AARF-17-533487 and BrightFocus Foundation fellowship A2015016F to SDG. AAV preparation was done with the help of Kazu Oka and the BCM Gene Vector Core. Slide scanning was done with the help of Cecilia Ljungberg at the BCM RNA In Situ Hybridization Core, supported by a Shared Instrumentation grant from the NIH (1S10OD016167) and the NIH IDDRC grant U54HD083092 from the Eunice Kennedy Shriver National Institute Of Child Health & Human Development. Confocal imaging was done with the help of Jason Kirk at the BCM Optical Imaging and Vital Microscopy Core.

## Contributions

SDG, RZ, and JLJ conceived the study. RZ, SDG, CAW, GAP, AKS, and ML performed experiments, RZ, SDG, CAW, GAP, MHL, AS, and MC analyzed data, KWP contributed reagents; JLJ wrote the manuscript with input from CAW, GAP, and SDG.

## Author information

The authors declare no competing financial interests.

---

**Supplemental Figure 1.**
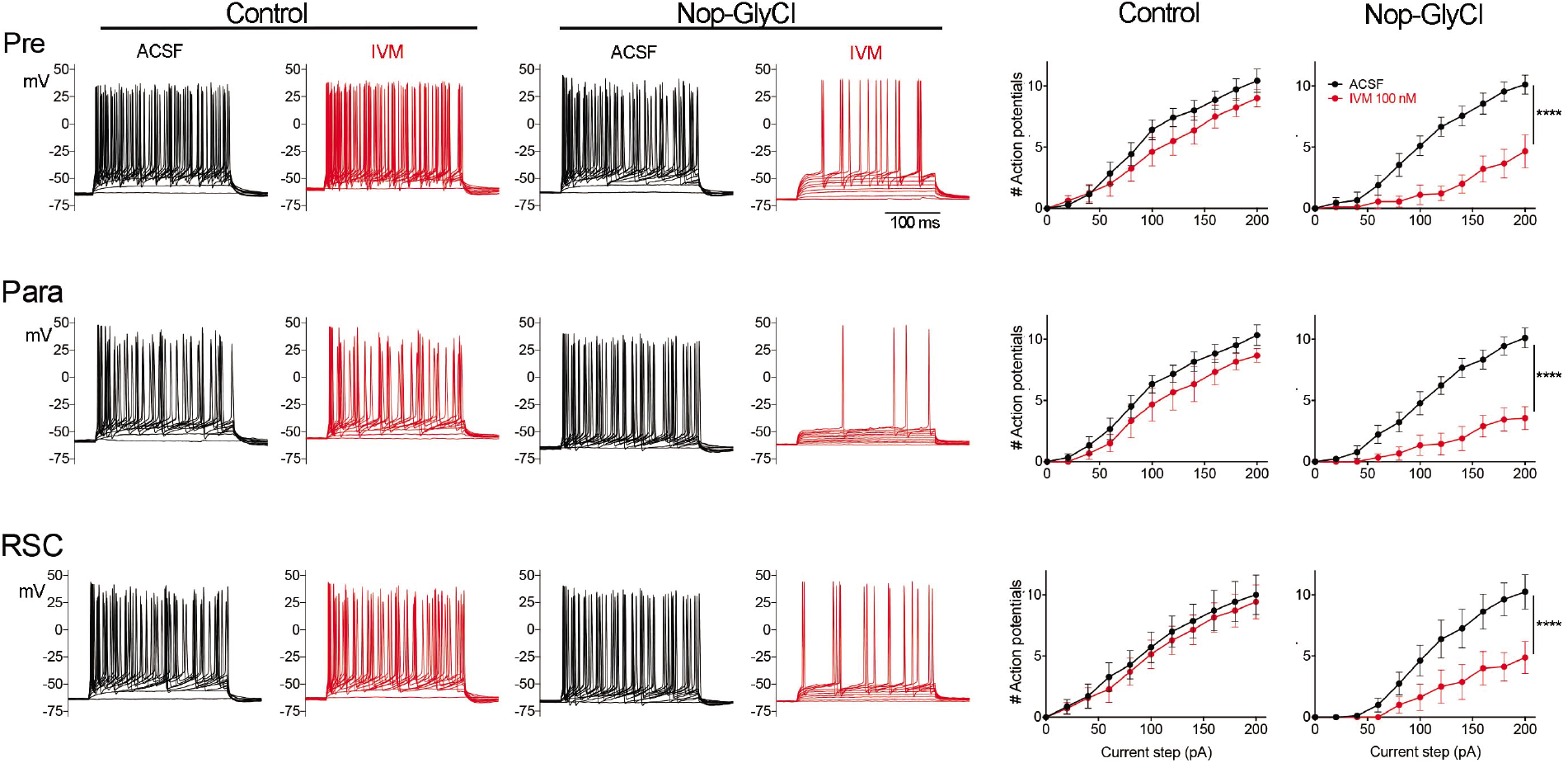
Brain regions outside EC express GlyCl in the Nop-GlyCl model and are electrically suppressed by IVM. Left panels show example traces from whole-cell current-clamp recordings in presubiculum (top row), parasubiculum (middle row), and retrosplenial cortex (bottom row). Action potential firing in response to stepped current injection is unaffected by application of 100 nM IVM in control mice (Nop-tTA single transgenic), but is decreased in cells from Nop-GlyCl bigenic animals. Summary graphs on right show action potential number as a function of current step (mean ± SEM). Graphs for Nop-GlyCl animals are identical to those shown in Figure 2 and are repeated here for comparison with controls. Data expressed as mean ± SEM, two-way repeated measure ANOVA (p value shown for interaction of current x genotype, ****p<0.0001). Controls: pre n=7 cells from 7 slices (3 mice); para n=6 cells from 6 slices (3 mice); RSC n=7 cells from 7 slices (5 mice).

**Supplemental Figure 2.**
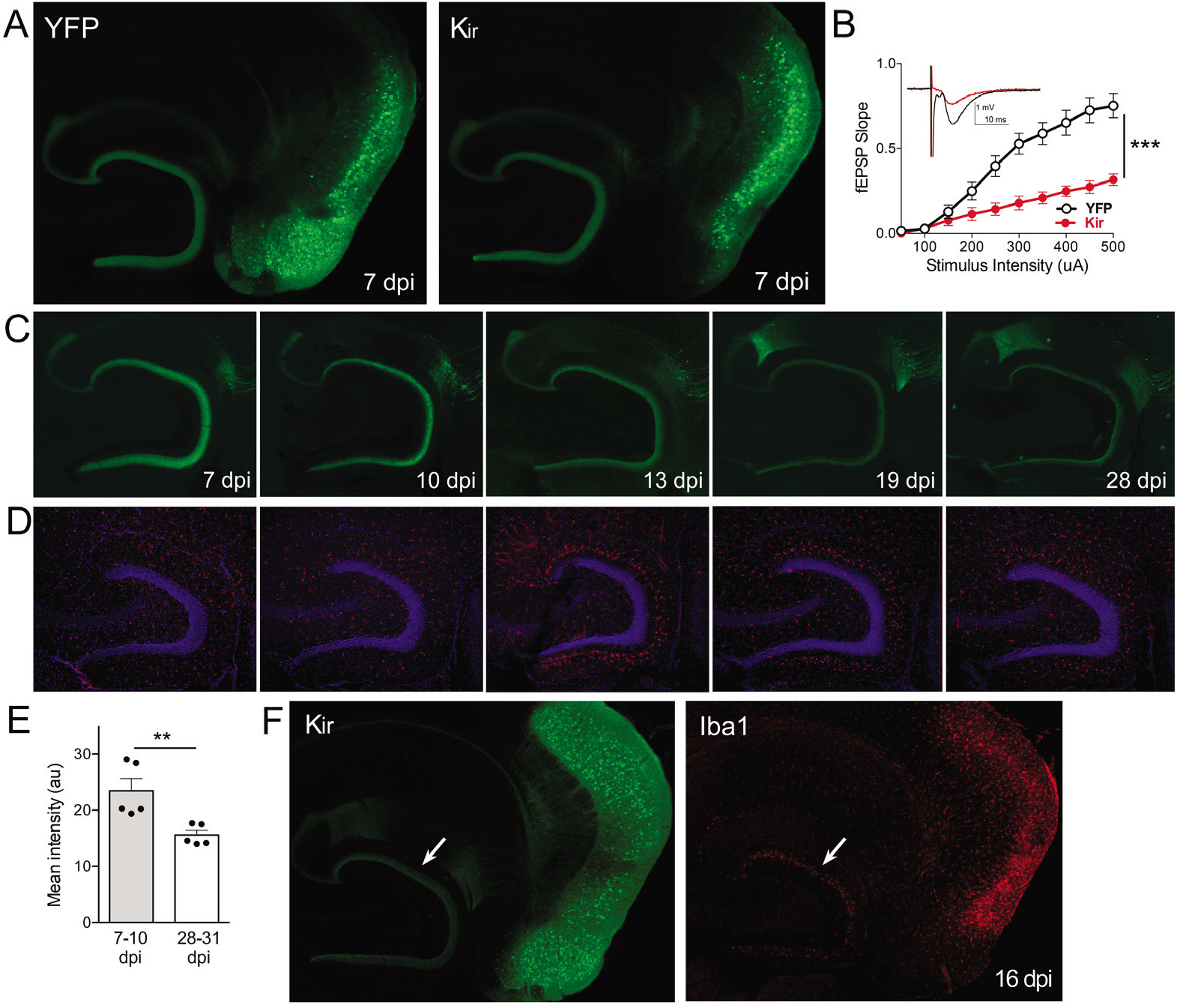
Gradual loss of DG labeling upon EC neuronal silencing with Kir. A. AAV was used to express either YFP or YFP plus Kir in entorhinal neurons via stereotaxic intracranial injection of adult Nop-tTA mice. By 7 dpi, labeled EC2 axons terminating in the DG can be clearly seen by either immuno- (shown) or native fluorescence in both YFP and Kir-YFP animals. B. Virally-delivered Kir effectively diminishes synaptic transmission into the DG. Data expressed as mean ± SEM, two-way repeated measure ANOVA (p value for genotype shown, ***p<0.001). n=15 slices from 3 mice (YFP), 10 slices from 3 mice (Kir), harvested 7-9 dpi. Note that YFP data shown here is identical to Figure 3B as the same control group was used for both experiments. C. Over the course of several weeks following injection, virally-delivered Kir leads to diminished labeling of perforant path axons. This phenotype is consistent with axonal loss, but lacked the axonal retraction bulbs of GlyCl. Images show native fluorescence. D. Iba1 immunostaining reveals hypertrophic microglia in the DG concurrent with loss of EC innervation. E. The intensity of YFP immunostaining in the DG decreased between the first and fourth weeks after viral injection of Kir. Graph data expressed as average values for each animal ± SEM, Student’s t-test. **p<0.01 F. Example of the microglial response to Kir expression at 16 dpi. YFP (left) and Iba1 (right) were co-immunostained to reveal hypertrophic microglia neighboring the silenced EC2 soma and their projections in DG (arrow). 7 dpi n=4 YFP, 4 Kir; 10 dpi n=3 YFP, 1 Kir; 12-13 dpi n=5 YFP, 3 Kir; 16-19 dpi n=10 YFP, 5 Kir; 22-28 dpi n=2 YFP, 5 Kir.

**Supplemental Figure 3.**
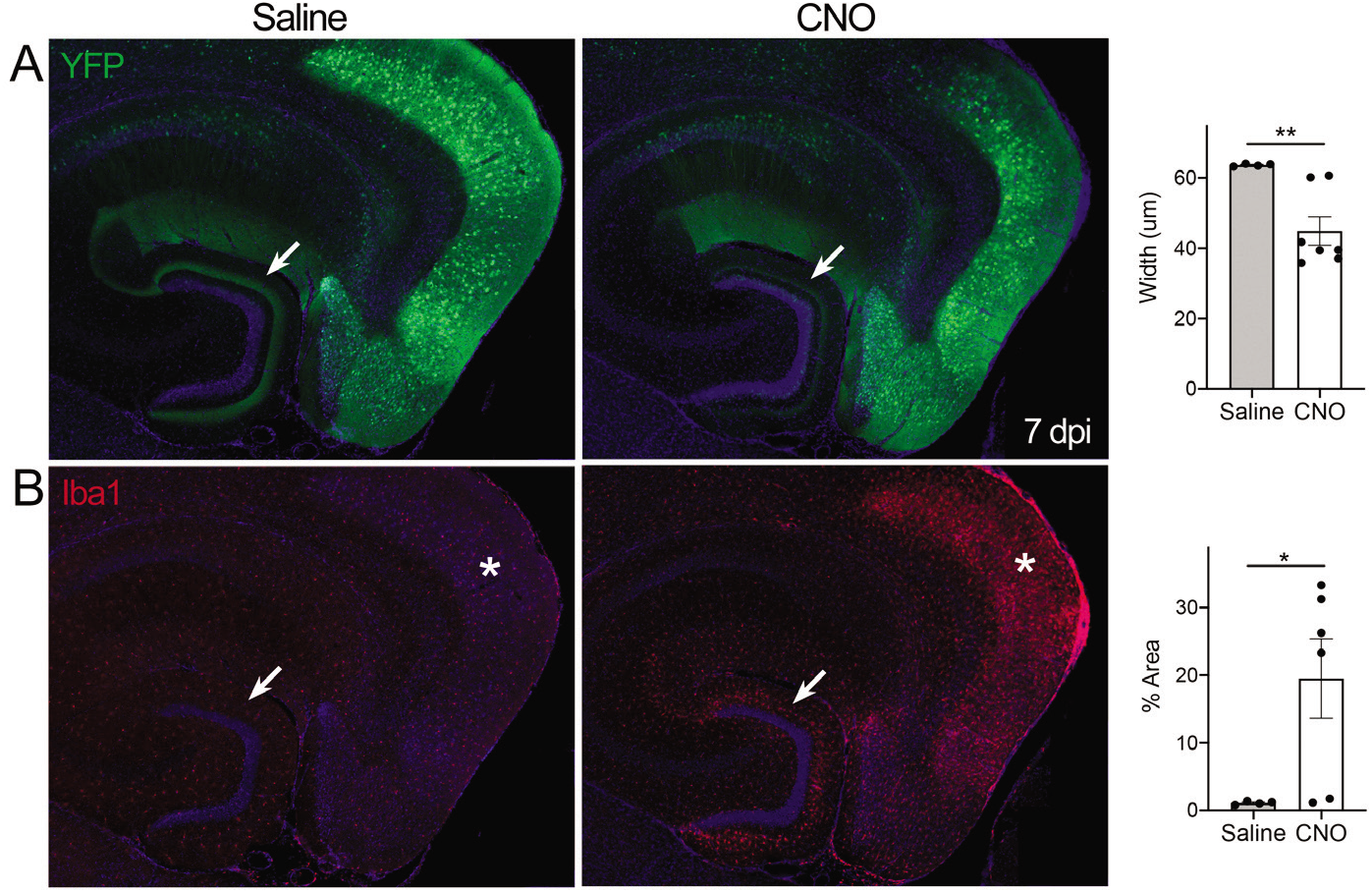
Silencing with hM4Di elicits axonal loss and microglial activation. A. AAV was used to express hM4Di plus YFP in entorhinal neurons via P0 intraventricular injection of Nop-tTA mice. Once the animals reached adulthood, they were implanted with mini-osmotic pumps to chronically deliver either CNO or saline for 3 days. 7 days after initial CNO exposure, treated animals have a thinner band of fluorescent labeling in the DG. Graph on right shows the reduction in width of YFP immunolabeling at the crest of the DG in CNO-treated mice. B. Iba1 immunostaining revealed the appearance of microglial activation in the DG and EC of CNO-treated animals but not saline controls. Graph on right shows the increase in % area occupied by Iba1 in the EC of CNO-treated mice. Graph data expressed as average values for each animal ± SEM, Student’s t-test with Welch’s correction. n=4 saline, 6 CNO. *p<0.05, ***p<0.001, ****p<0.0001.

**Supplemental Figure 4.**
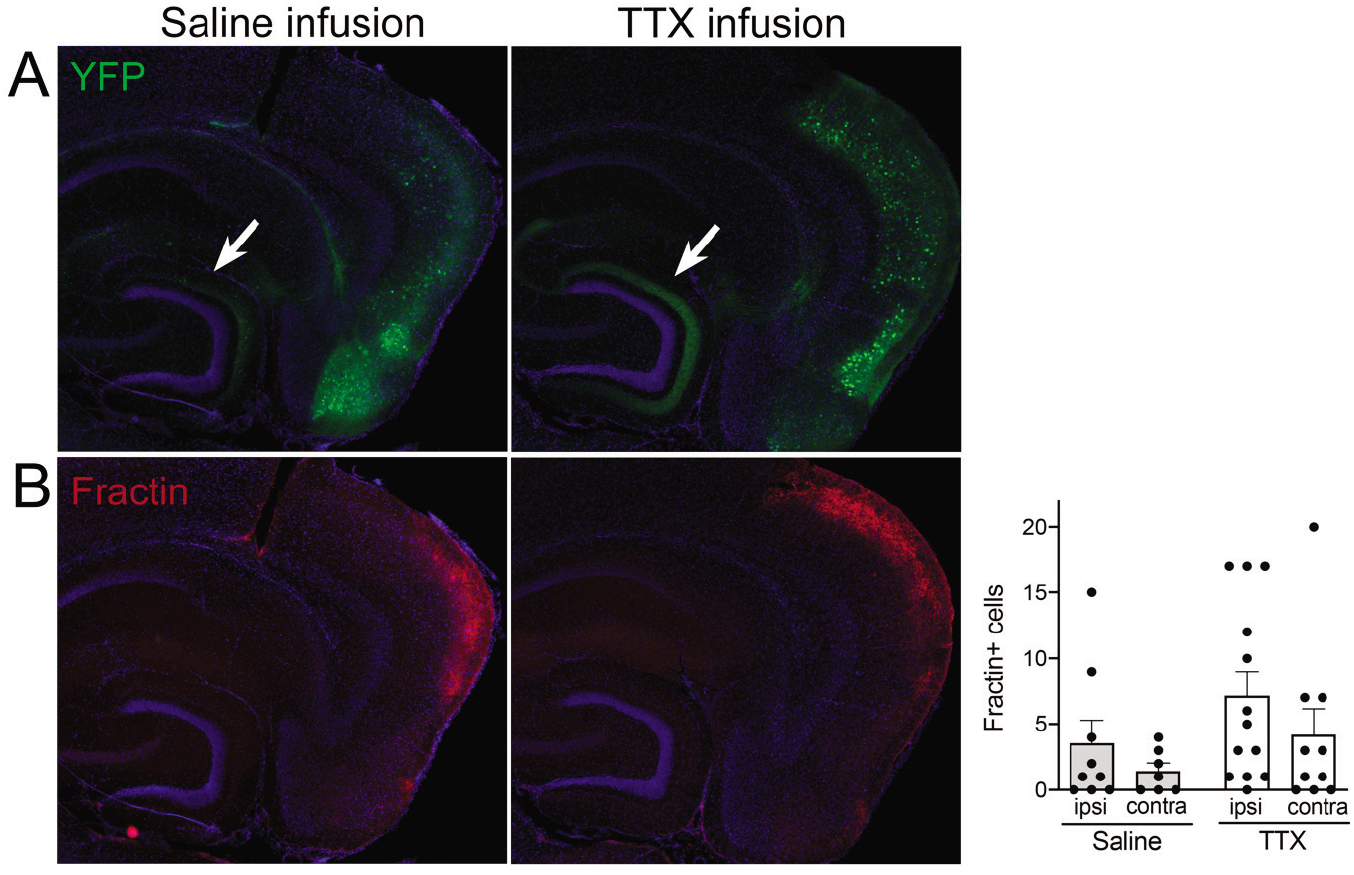
Appearance of fractin due to TeTX silencing is not changed by TTX infusion. Mice expressing TeTX+YFP in EC neurons were treated by local infusion of TTX or saline. Infusion began 3 days after viral injection and continued until mice were harvested 7 days later (10 dpi). A. Images show native YFP fluorescence in TeTX mice following saline (left) or TTX (right) infusion. Note the retention of DG innervation in mice treated with TTX compared to the loss of innervation with saline infusion (arrows). B. Although axonal YFP labeling was protected by TTX, the number of fractin+ cells in the EC at this time point did not differ between conditions. The location of fractin labeling appeared to shift more dorsal and lateral in TTX-infused mice, and the extent of neurite labeling (as opposed to soma) was greater with 3/9 (saline) vs 8/13 (TTX) mice harboring some EC fractin. Graph data expressed as average total fractin+ cells per animal ± SEM, one-way ANOVA with Tukey’s post-hoc test (missed injection target for some hemispheres precluded 2-way ANOVA). All comparisons n.s. n=7-9 (Saline infusion), 10-13 (TTX infusion)

## References

Armbruster, B.N., Li, X., Pausch, M.H., Herlitze, S., and Roth, B.L. (2007). Evolving the lock to fit the key to create a family of G protein-coupled receptors potently activated by an inert ligand. Proc Natl Acad Sci U S A 104, 5163–5168.

Assali, A., Gaspar, P., and Rebsam, A. (2014). Activity dependent mechanisms of visual map formation--from retinal waves to molecular regulators. Semin Cell Dev Biol 35, 136–146.

Bavelier, D., Levi, D.M., Li, R.W., Dan, Y., and Hensch, T.K. (2010). Removing brakes on adult brain plasticity: from molecular to behavioral interventions. J Neurosci 30, 14964–14971.

Blakemore, C. (1991). Sensitive and vulnerable periods in the development of the visual system. Ciba Found Symp 156, 129–147; discussion 147-154.

Blanquie, O., Kilb, W., Sinning, A., and Luhmann, H.J. (2017). Homeostatic interplay between electrical activity and neuronal apoptosis in the developing neocortex. Neuroscience 358, 190–200.

Bothwell, M., and Giniger, E. (2000). Alzheimer’s disease: neurodevelopment converges with neurodegeneration. Cell 102, 271–273.

Braak, H., and Braak, E. (1991). Neuropathological stageing of Alzheimer-related changes. Acta Neuropathol (Berl) 82, 239–259.

Braak, H., and Braak, E. (1995). Staging of Alzheimer’s disease-related neurofibrillary changes. Neurobiol Aging 16, 271–278; discussion 278-284.

Cirrito, J.R., Yamada, K.A., Finn, M.B., Sloviter, R.S., Bales, K.R., May, P.C., Schoepp, D.D., Paul, S.M., Mennerick, S., and Holtzman, D.M. (2005). Synaptic activity regulates interstitial fluid amyloid-beta levels in vivo. Neuron 48, 913–922.

Crain, B.J., and Burger, P.C. (1988). The laminar distribution of neuritic plaques in the fascia dentata of patients with Alzheimer’s disease. Acta Neuropathol 76, 87–93.

Dancause, N. (2006). Neurophysiological and anatomical plasticity in the adult sensorimotor cortex. Reviews in the neurosciences 17, 561–580.

de Villers-Sidani, E., and Merzenich, M.M. (2011). Lifelong plasticity in the rat auditory cortex: basic mechanisms and role of sensory experience. Progress in brain research 191, 119–131.

Fox, K., and Wong, R.O. (2005). A comparison of experience-dependent plasticity in the visual and somatosensory systems. Neuron 48, 465–477.

Franklin, K.B.J., and Paxinos, G. (2008). The mouse brain in stereotaxic coordinates, 3rd edn (San Diego, CA: Acadmic Press).

Geddes, J.W., Anderson, K.J., and Cotman, C.W. (1986). Senile plaques as aberrant sprout-stimulating structures. Exp Neurol 94, 767–776.

Gomez-Isla, T., Price, J.L., McKeel, D.W., Jr., Morris, J.C., Growdon, J.H., and Hyman, B.T. (1996). Profound loss of layer II entorhinal cortex neurons occurs in very mild Alzheimer’s disease. J Neurosci 16, 4491–4500.

Gonzalez-Rodriguez, P., Zampese, E., and Surmeier, D.J. (2020). Selective neuronal vulnerability in Parkinson’s disease. Progress in brain research 252, 61–89.

Harris, S.S., Wolf, F., De Strooper, B., and Busche, M.A. (2020). Tipping the Scales: Peptide-Dependent Dysregulation of Neural Circuit Dynamics in Alzheimer’s Disease. Neuron 107, 417–435.

Hensch, T.K., and Fagiolini, M. (2005). Excitatory-inhibitory balance and critical period plasticity in developing visual cortex. Progress in brain research 147, 115–124.

Heys, J.G., Wu, Z., Allegra Mascaro, A.L., and Dombeck, D.A. (2020). Inactivation of the Medial Entorhinal Cortex Selectively Disrupts Learning of Interval Timing. Cell reports 32, 108163.

Hill, C.S., Coleman, M.P., and Menon, D.K. (2016). Traumatic Axonal Injury: Mechanisms and Translational Opportunities. Trends in neurosciences 39, 311–324.

Hollville, E., and Deshmukh, M. (2018). Physiological functions of non-apoptotic caspase activity in the nervous system. Semin Cell Dev Biol 82, 127–136.

Hubener, M., and Bonhoeffer, T. (2014). Neuronal plasticity: beyond the critical period. Cell 159, 727–737.

Huichalaf, C.H., Al-Ramahi, I., Park, K.W., Grunke, S.D., Lu, N., de Haro, M., El-Zein, K., Gallego-Flores, T., Perez, A.M., Jung, S.Y., et al. (2019). Cross-species genetic screens to identify kinase targets for APP reduction in Alzheimer’s disease. Hum Mol Genet 28, 2014–2029.

Humeau, Y., Doussau, F., Grant, N.J., and Poulain, B. (2000). How botulinum and tetanus neurotoxins block neurotransmitter release. Biochimie 82, 427–446.

Hyman, B.T., Van Hoesen, G.W., Damasio, A.R., and Barnes, C.L. (1984). Alzheimer’s disease: cell-specific pathology isolates the hippocampal formation. Science 225, 1168–1170.

Hyman, B.T., Van Hoesen, G.W., Kromer, L.J., and Damasio, A.R. (1986). Perforant pathway changes and the memory impairment of Alzheimer’s disease. Ann Neurol 20, 472–481.

Kanter, B.R., Lykken, C.M., Avesar, D., Weible, A., Dickinson, J., Dunn, B., Borgesius, N.Z., Roudi, Y., and Kentros, C.G. (2017). A Novel Mechanism for the Grid-to-Place Cell Transformation Revealed by Transgenic Depolarization of Medial Entorhinal Cortex Layer II. Neuron 93, 1480–1492 e1486.

Kim, J.Y., Grunke, S.D., and Jankowsky, J.L. (2016). Widespread Neuronal Transduction of the Rodent CNS via Neonatal Viral Injection. Methods Mol Biol 1382, 239–250.

Kim, J.Y., Grunke, S.D., Levites, Y., Golde, T.E., and Jankowsky, J.L. (2014). Intracerebroventricular viral injection of the neonatal mouse brain for persistent and widespread neuronal transduction. J Vis Exp, 51863.

Kordower, J.H., Chu, Y., Stebbins, G.T., DeKosky, S.T., Cochran, E.J., Bennett, D., and Mufson, E.J. (2001). Loss and atrophy of layer II entorhinal cortex neurons in elderly people with mild cognitive impairment. Ann Neurol 49, 202–213.

Lerchner, W., Xiao, C., Nashmi, R., Slimko, E.M., van Trigt, L., Lester, H.A., and Anderson, D.J. (2007). Reversible silencing of neuronal excitability in behaving mice by a genetically targeted, ivermectin-gated Cl-channel. Neuron 54, 35–49.

Leung, C., Cao, F., Nguyen, R., Joshi, K., Aqrabawi, A.J., Xia, S., Cortez, M.A., Snead, O.C., 3rd, Kim, J.C., and Jia, Z. (2018). Activation of Entorhinal Cortical Projections to the Dentate Gyrus Underlies Social Memory Retrieval. Cell reports 23, 2379–2391.

Li, M.H., Suchland, K.L., and Ingram, S.L. (2015). GABAergic transmission and enhanced modulation by opioids and endocannabinoids in adult rat rostral ventromedial medulla. The Journal of physiology 593, 217–230.

Li, M.H., Suchland, K.L., and Ingram, S.L. (2017). Compensatory Activation of Cannabinoid CB2 Receptor Inhibition of GABA Release in the Rostral Ventromedial Medulla in Inflammatory Pain. J Neurosci 37, 626–636.

Lopez, C.M., Pelkey, K.A., Chittajallu, R., Nakashiba, T., Toth, K., Tonegawa, S., and McBain, C.J. (2012). Competition from newborn granule cells does not drive axonal retraction of silenced old granule cells in the adult hippocampus. Front Neural Circuits 6, 85.

Lynagh, T., and Lynch, J.W. (2010). An improved ivermectin-activated chloride channel receptor for inhibiting electrical activity in defined neuronal populations. J Biol Chem 285, 14890–14897.

Lynagh, T., and Lynch, J.W. (2012). Molecular mechanisms of Cys-loop ion channel receptor modulation by ivermectin. Frontiers in molecular neuroscience 5, 60.

Martin, J.H. (2016). Harnessing neural activity to promote repair of the damaged corticospinal system after spinal cord injury. Neural Regen Res 11, 1389–1391.

Mayford, M., Baranes, D., Podsypanina, K., and Kandel, E.R. (1996). The 3’-untranslated region of CaMKII alpha is a cis-acting signal for the localization and translation of mRNA in dendrites. Proc Natl Acad Sci U S A 93, 13250–13255.

Miao, C., Cao, Q., Ito, H.T., Yamahachi, H., Witter, M.P., Moser, M.B., and Moser, E.I. (2015). Hippocampal Remapping after Partial Inactivation of the Medial Entorhinal Cortex. Neuron 88, 590–603.

Mukherjee, A., and Williams, D.W. (2017). More alive than dead: non-apoptotic roles for caspases in neuronal development, plasticity and disease. Cell Death Differ 24, 1411–1421.

Murase, S. (2014). A new model for developmental neuronal death and excitatory/inhibitory balance in hippocampus. Molecular neurobiology 49, 316–325.

Murase, S., Owens, D.F., and McKay, R.D. (2011). In the newborn hippocampus, neurotrophin-dependent survival requires spontaneous activity and integrin signaling. J Neurosci 31, 7791–7800.

Nahmani, M., and Turrigiano, G.G. (2014). Adult cortical plasticity following injury: Recapitulation of critical period mechanisms? Neuroscience 283, 4–16.

Nakashiba, T., Cushman, J.D., Pelkey, K.A., Renaudineau, S., Buhl, D.L., McHugh, T.J., Rodriguez Barrera, V., Chittajallu, R., Iwamoto, K.S., McBain, C.J., et al. (2012). Young dentate granule cells mediate pattern separation, whereas old granule cells facilitate pattern completion. Cell 149, 188–201.

Nakashiba, T., Young, J.Z., McHugh, T.J., Buhl, D.L., and Tonegawa, S. (2008). Transgenic inhibition of synaptic transmission reveals role of CA3 output in hippocampal learning. Science 319, 1260–1264.

Neukomm, L.J., and Freeman, M.R. (2014). Diverse cellular and molecular modes of axon degeneration. Trends Cell Biol 24, 515–523.

Nguyen, M.D., Mushynski, W.E., and Julien, J.P. (2002). Cycling at the interface between neurodevelopment and neurodegeneration. Cell Death Differ 9, 1294–1306.

Ormond, J., and McNaughton, B.L. (2015). Place field expansion after focal MEC inactivations is consistent with loss of Fourier components and path integrator gain reduction. Proc Natl Acad Sci U S A 112, 4116–4121.

Palop, J.J., and Mucke, L. (2016). Network abnormalities and interneuron dysfunction in Alzheimer disease. Nat Rev Neurosci 17, 777–792.

Park, K.W., Wood, C.A., Li, J., Taylor, B.C., Oh, S., Young, N.L., and Jankowsky, J.L. (2021). Gene therapy using Abeta variants for amyloid reduction. Molecular therapy : the journal of the American Society of Gene Therapy.

Portera-Cailliau, C., Weimer, R.M., De Paola, V., Caroni, P., and Svoboda, K. (2005). Diverse modes of axon elaboration in the developing neocortex. PLoS Biol 3, e272.

Qin, H., Fu, L., Hu, B., Liao, X., Lu, J., He, W., Liang, S., Zhang, K., Li, R., Yao, J., et al. (2018). A Visual-Cue-Dependent Memory Circuit for Place Navigation. Neuron 99, 47–55 e44.

Reh, R.K., Dias, B.G., Nelson, C.A., 3rd, Kaufer, D., Werker, J.F., Kolb, B., Levine, J.D., and Hensch, T.K. (2020). Critical period regulation across multiple timescales. Proc Natl Acad Sci U S A 117, 23242–23251.

Riccomagno, M.M., and Kolodkin, A.L. (2015). Sculpting neural circuits by axon and dendrite pruning. Annual review of cell and developmental biology 31, 779–805.

Robinson, N.T.M., Priestley, J.B., Rueckemann, J.W., Garcia, A.D., Smeglin, V.A., Marino, F.A., and Eichenbaum, H. (2017). Medial Entorhinal Cortex Selectively Supports Temporal Coding by Hippocampal Neurons. Neuron 94, 677–688 e676.

Rodriguez, G.A., Barrett, G.M., Duff, K.E., and Hussaini, S.A. (2020). Chemogenetic attenuation of neuronal activity in the entorhinal cortex reduces Abeta and tau pathology in the hippocampus. PLoS Biol 18, e3000851.

Rowland, D.C., Obenhaus, H.A., Skytoen, E.R., Zhang, Q., Kentros, C.G., Moser, E.I., and Moser, M.B. (2018). Functional properties of stellate cells in medial entorhinal cortex layer II. eLife 7.

Rueckemann, J.W., DiMauro, A.J., Rangel, L.M., Han, X., Boyden, E.S., and Eichenbaum, H. (2016). Transient optogenetic inactivation of the medial entorhinal cortex biases the active population of hippocampal neurons. Hippocampus 26, 246–260.

Salimi, I., Friel, K.M., and Martin, J.H. (2008). Pyramidal tract stimulation restores normal corticospinal tract connections and visuomotor skill after early postnatal motor cortex activity blockade. J Neurosci 28, 7426–7434.

Schiavo, G., Benfenati, F., Poulain, B., Rossetto, O., Polverino de Laureto, P., DasGupta, B.R., and Montecucco, C. (1992). Tetanus and botulinum-B neurotoxins block neurotransmitter release by proteolytic cleavage of synaptobrevin. Nature 359, 832–835.

Schulz, R., Vogel, T., Mashima, T., Tsuruo, T., and Krieglstein, K. (2009). Involvement of Fractin in TGF-beta-induced apoptosis in oligodendroglial progenitor cells. Glia 57, 1619–1629.

Spolidoro, M., Sale, A., Berardi, N., and Maffei, L. (2009). Plasticity in the adult brain: lessons from the visual system. Experimental brain research Experimentelle Hirnforschung Experimentation cerebrale 192, 335–341.

Ting, J.T., Daigle, T.L., Chen, Q., and Feng, G. (2014). Acute brain slice methods for adult and aging animals: application of targeted patch clamp analysis and optogenetics. Methods Mol Biol 1183, 221–242.

Van Hoesen, G.W., Hyman, B.T., and Damasio, A.R. (1986). Cell-specific pathology in neural systems of the temporal lobe in Alzheimer’s disease. Progress in brain research 70, 321–335.

Van Hoesen, G.W., Hyman, B.T., and Damasio, A.R. (1991). Entorhinal cortex pathology in Alzheimer’s disease. Hippocampus 1, 1–8.

van Wijngaarden, J.B., Babl, S.S., and Ito, H.T. (2020). Entorhinal-retrosplenial circuits for allocentric-egocentric transformation of boundary coding. eLife 9.

Wang, C., and Holtzman, D.M. (2020). Bidirectional relationship between sleep and Alzheimer’s disease: role of amyloid, tau, and other factors. Neuropsychopharmacology 45, 104–120.

Weir, G.A., Middleton, S.J., Clark, A.J., Daniel, T., Khovanov, N., McMahon, S.B., and Bennett, D.L. (2017). Using an engineered glutamate-gated chloride channel to silence sensory neurons and treat neuropathic pain at the source. Brain 140, 2570–2585.

Wong, F.K., and Marin, O. (2019). Developmental Cell Death in the Cerebral Cortex. Annual review of cell and developmental biology 35, 523–542.

Wu, J.W., Hussaini, S.A., Bastille, I.M., Rodriguez, G.A., Mrejeru, A., Rilett, K., Sanders, D.W., Cook, C., Fu, H., Boonen, R.A., et al. (2016). Neuronal activity enhances tau propagation and tau pathology in vivo. Nat Neurosci 19, 1085–1092.

Xue, M., Atallah, B.V., and Scanziani, M. (2014). Equalizing excitation-inhibition ratios across visual cortical neurons. Nature 511, 596–600.

Yamada, K., Holth, J.K., Liao, F., Stewart, F.R., Mahan, T.E., Jiang, H., Cirrito, J.R., Patel, T.K., Hochgrafe, K., Mandelkow, E.M., and Holtzman, D.M. (2014). Neuronal activity regulates extracellular tau in vivo. The Journal of experimental medicine 211, 387–393.

Yamamoto, K., Tanei, Z.I., Hashimoto, T., Wakabayashi, T., Okuno, H., Naka, Y., Yizhar, O., Fenno, L.E., Fukayama, M., Bito, H., et al. (2015). Chronic optogenetic activation augments abeta pathology in a mouse model of Alzheimer disease. Cell reports 11, 859–865.

Yang, F., Sun, X., Beech, W., Teter, B., Wu, S., Sigel, J., Vinters, H.V., Frautschy, S.A., and Cole, G.M. (1998). Antibody to caspase-cleaved actin detects apoptosis in differentiated neuroblastoma and plaque-associated neurons and microglia in Alzheimer’s disease. Am J Pathol 152, 379–389.

Yasuda, M., Johnson-Venkatesh, E.M., Zhang, H., Parent, J.M., Sutton, M.A., and Umemori, H. (2011). Multiple forms of activity-dependent competition refine hippocampal circuits in vivo. Neuron 70, 1128–1142.

Yasuda, M., and Mayford, M.R. (2006). CaMKII activation in the entorhinal cortex disrupts previously encoded spatial memory. Neuron 50, 309–318.

Yetman, M.J., Lillehaug, S., Bjaalie, J.G., Leergaard, T.B., and Jankowsky, J.L. (2015). Transgene expression in the Nop-tTA driver line is not inherently restricted to the entorhinal cortex. Brain structure & function.

Yu, C.R., Power, J., Barnea, G., O’Donnell, S., Brown, H.E., Osborne, J., Axel, R., and Gogos, J.A. (2004). Spontaneous neural activity is required for the establishment and maintenance of the olfactory sensory map. Neuron 42, 553–566.

Yuan, P., and Grutzendler, J. (2016). Attenuation of beta-Amyloid Deposition and Neurotoxicity by Chemogenetic Modulation of Neural Activity. J Neurosci 36, 632–641.

Yun, S., Reynolds, R.P., Petrof, I., White, A., Rivera, P.D., Segev, A., Gibson, A.D., Suarez, M., DeSalle, M.J., Ito, N., et al. (2018). Stimulation of entorhinal cortex-dentate gyrus circuitry is antidepressive. Nat Med 24, 658–666.

Zhang, S.J., Ye, J., Couey, J.J., Witter, M., Moser, E.I., and Moser, M.B. (2014). Functional connectivity of the entorhinal-hippocampal space circuit. Philosophical transactions of the Royal Society of London Series B, Biological sciences 369, 20120516.

Zhang, S.J., Ye, J., Miao, C., Tsao, A., Cerniauskas, I., Ledergerber, D., Moser, M.B., and Moser, E.I. (2013). Optogenetic dissection of entorhinal-hippocampal functional connectivity. Science 340, 1232627.

Zhao, H., and Reed, R.R. (2001). X inactivation of the OCNC1 channel gene reveals a role for activity-dependent competition in the olfactory system. Cell 104, 651–660.

Zhao, R., Grunke, S.D., Keralapurath, M.M., Yetman, M.J., Lam, A., Lee, T.C., Sousounis, K., Jiang, Y., Swing, D.A., Tessarollo, L., et al. (2016). Impaired Recall of Positional Memory following Chemogenetic Disruption of Place Field Stability. Cell reports 16, 793–804.

Zhu, H., Pleil, K.E., Urban, D.J., Moy, S.S., Kash, T.L., and Roth, B.L. (2014). Chemogenetic inactivation of ventral hippocampal glutamatergic neurons disrupts consolidation of contextual fear memory. Neuropsychopharmacology 39, 1880–1892.

